# Protein domain-based structural interfaces help interpret biologically-relevant interactions in the human interaction network

**DOI:** 10.1101/2020.03.14.992149

**Authors:** Krishna Praneeth Kilambi, Qifang Xu, Guruharsha Kuthethur Gururaj, Kejie Li, Spyros Artavanis-Tsakonas, Roland L. Dunbrack, Andreas Lehmann

## Abstract

A high-quality map of the human protein–protein interaction (PPI) network can help us better understand complex genotype–phenotype relationships. Each edge between two interacting proteins supported through an interface in a three-dimensional (3D) structure of a protein complex adds credibility to the biological relevance of the interaction. Such structure-supported interactions would augment an interaction map primarily built using high-throughput cell-based biophysical methods. Here, we integrate structural information with the human PPI network to build the structure-supported human interactome, a subnetwork of PPI between proteins that contain domains or regions known to form interfaces in the 3D structures of protein complexes. We expand the coverage of our structure-supported human interactome by using Pfam-based domain definitions, whereby we include homologous interactions if a human complex structure is unavailable. The structure-supported interactome predicts one-eighth of the total network PPI to interact through domain–domain interfaces. It identifies with higher resolution the interacting subunits in multi-protein complexes and enables us to characterize functional and disease-relevant neighborhoods in the network map with higher accuracy, allowing for structural insights into disease-associated genes and pathways. We expand the structural coverage beyond domain–domain interfaces by identifying the most common non-enzymatic peptide-binding domains with structural support. Adding these interactions between protein domains on one side and peptide regions on the other approximately doubles the number of structure-supported PPI. The human structure-supported interactome is a resource to prioritize investigations of smaller-scale context-specific experimental PPI neighborhoods of biological or clinical significance.

**Short abstract:** A high-quality map of the human protein–protein interaction (PPI) network can help us better understand genotype–phenotype relationships. Each edge between two interacting proteins supported through an interface in a three-dimensional structure of a protein complex adds credibility to the biological relevance of the interaction aiding experimental prioritization. Here, we integrate structural information with the human interactome to build the structure-supported human interactome, a subnetwork of PPI between proteins that contain domains or regions known to form interfaces in the structures of protein complexes. The structure-supported interactome predicts one-eighth of the total PPI to interact through domain–domain interfaces. It identifies with higher resolution the interacting subunits in multi-protein complexes and enables us to structurally characterize functional, disease-relevant network neighborhoods. We also expand the structural coverage by identifying PPI between non-enzymatic peptide-binding domains on one side and peptide regions on the other, thereby doubling the number of structure-supported PPI.

## Introduction

Protein–protein interactions (PPI) are fundamental to all cellular processes. Comprehensive large-scale mapping of the human PPI has been the topic of several extensive studies.^1–4^ Current high-throughput efforts focused on generating the human interactome broadly rely on either yeast two-hybrid (Y2H) or affinity purification–mass spectrometry (AP–MS) based biophysical methods. These studies have produced datasets offering an unprecedented view of the functional organization of protein interaction modules in the human network. BioPlex,^2^ the largest of these efforts to date, includes 10,961 proteins covering over half of the human proteome. However, these studies rely on statistical methods to eliminate false-positive PPI, and they miss many transient interactions and are blind to potential posttranslational modifications hypothesized to dramatically increase the diversity of the human proteome.^5,6^ For these reasons, the overlap between the different human PPI datasets is still small, and a large fraction of protein interactions likely remain to be discovered.^7,8^ Despite these challenges, researchers have used the existing human PPI network to identify disease modules and demonstrated that network-based approaches to studying human disease can partly overcome the incompleteness of the interactome.^9^ Structure-based interfaces constitute a high-quality dataset that can be exploited to enable a higher-resolution picture of the protein–protein interactions (PPI) in the network. In this study, we integrate domain–domain and domain–peptide interfaces from 3D structures of protein–protein complexes in the Protein Data Bank (PDB)^10^ with the human interactome and demonstrate that structural data can be used to extract more detailed information from the PPI network.

Structural data were used previously to model existing PPI, and to predict new PPI in interaction networks.^11^ Yu and co-workers integrated structural data with binary human PPI to construct the hSIN network^12^ with over 4200 interactions. They mapped nearly 22,000 disease-associated mutations to the network proteins, and later expanded the mutation mapping to more binary PPI.^13^ Aloy and co-workers developed Interactome3D^14^ based on the 3did^15^ database, a resource containing structural details for over 12,000 binary PPI from interactomes in eight model organisms, and tools to build 3D models for new PPI based on structural templates. They also later mapped disease-related missense mutations to the human interactome.^16^ Honig and co-workers exploited the predictive capabilities of structural data to build PrePPI, a computational proteome-scale map of the human interaction network.^17^ Besides structural modeling, PrePPI also combines data from gene ontology, orthology, and phylogenetic and protein expression profiles to predict over 1.3 million probable PPI.^18^ However, these studies mostly ignore the interactions mediated by proteins with intrinsically disordered regions (IDRs), often involving post-translational modifications, that are prevalent in cellular regulation and signaling^19^ and are estimated to compose up to 40% of the interactome.^20^ Although PrePPI was later expanded to include protein–peptide interactions,^21^ its primary aim was prediction of novel PPI rather than offering structural insights on experimentally derived PPI.

We previously described development of the Protein Common Interfaces Database (ProtCID^22^), which contains structural clusters of similar homo-and heterodimeric interfaces observed in multiple crystal forms (CFs) of protein 3D structures deposited in the PDB. Domain–domain interfaces that repeatedly occur in multiple CFs, and that belong to homologous proteins across different species are often biologically relevant, i.e., these interfaces likely occur *in vivo*.^23^ ProtCID facilitates the systematic exploration of biological and secondary interfaces. It contains over 14,000 unique domain–domain interfaces and has also recently been expanded to include domain–peptide, domain–ligand, and domain– DNA/RNA interfaces.^24^ Here, we present a domain-based approach using biological interfaces from the ProtCID database to identify structure-supported PPI within the human interaction network. First, we methodically assign Pfam (protein family) domains to the human proteome using an updated version of the multi-step greedy algorithm we developed for our previous study.^25^ Second, we use structure-supported biological domain–domain interfaces from ProtCID to identify the structure-supported subset of PPI within a larger model human interaction network obtained by merging interactions from three high-quality networks.^1,2^ Third, we map known disease-associated mutations in proteins to their individual domains and demonstrate that structural interfaces offer a more detailed picture of the PPI affected by the disease variants. Finally, we identify the most widely-used structure-supported non-enzymatic peptide-binding domains in the human proteome and use these to expand the coverage of the structure-supported interactome by predicting interactions involving domain–peptide interfaces in the experimental PPI network.

## Results

### A new Pfam assignment method improves accuracy of repeat and split domain assignments

To identify the structure-supported PPIs within the network, we first assigned Pfam domains^26^ to the protein sequences in the human Uniprot Reference Proteome.^27^ These Pfam domain assignments enabled us to map the Pfam-based interfaces in ProtCID to the PPI that connect the proteins in the network. For example, a biological interface between two Pfam domains in ProtCID provides structural support for the biological relevance of an edge that connects two protein nodes in the PPI network, if each protein node contains one of the two interacting Pfam domains. To assign Pfams to the human Uniprot Reference Proteome, we modified a greedy algorithm we previously developed to assign Pfams to sequences of protein structures in the Protein Data Bank (PDB).^25^ The algorithm uses alignments of human sequences to Pfam hidden-Markov models (HMMs, which are statistical models of multiple sequence alignments of homologous protein domains) with the program HMMER and HMM-HMM alignments of HMMs of human sequences to the Pfam HMMs with HHsearch (Supplementary Fig. S1). Hits are assigned in order of their statistical significance; hits cannot be aligned to regions already covered by a Pfam earlier in the procedure. Often alignments to Pfam HMMs contain large unaligned regions of the query sequence flanked by aligned regions because of domains inserted into the human protein during evolution. We refer to these as “split” domains; the algorithm allows these inserted regions to be assigned Pfams in subsequent steps of the procedure. Pfams of repeat regions are assigned with less stringent E-value cutoffs if there are already instances of the same repeats assigned to the sequence with strong E-values.

We used the algorithm to assign 56,801 Pfam domains to 19,032 proteins in the human Uniprot Reference Proteome and compared our assignments (HuPfam) to those obtained from the online Pfam database (Pfam) for the 18,389 human proteins common to both data sets (Supplementary Table S1). Overall, HuPfam covers 11% more sequence residues and 12% more Pfam HMM residues compared to Pfam and captures 117 Pfams not observed even once in Pfam’s assignments (6198 vs. 6081 different Pfams).

Remarkably, HuPfam assigns 60% more repeat Pfams compared to Pfam as our algorithm considers weak Pfam repeats if there are other higher confidence repeat assignments of the same Pfam within the protein sequence. For example, HuPfam correctly identifies all 16 LRR repeats in Ribonuclease inhibitor (RINI) protein, according to the structure, while there are only two copies of the repeat assigned in Pfam (Fig. 1a). Similarly, the more sensitive HMM-HMM alignments capture the three TPR repeats in AH receptor-interacting protein (AIP), whereas no repeats are assigned by Pfam (Fig. 1b). In the case of split domains, the presence of large insertions often results in two alignments to the Pfam covering different parts of the protein sequence. HuPfam combines these split assignments into one domain, whereas in Pfam, it is either a larger assignment overlapping the insertion sequence, or two assignments, which may be confused with two independent copies of the Pfam. HuPfam identifies 1,000 such split domains across 963 human proteins. Figures 1c-d show two examples of split domains in Ubiquitin-like modifier-activating enzyme 1 (UBA1) and Beta-Ala-His dipeptidase (CNDP1). Pfam assigns a ThiF domain to residues 55-450 of UBA1, while HuPfam assigns it a single ThiF domain to residues 55-225 and 389-450. Additionally, HuPfam assigns an inserted E1_FCCH domain to residues 227-297 and an E1_4HB domain to 298-366 of UBA1. Similarly, HuPfam also correctly assigns the M20_dimer domain - itself a split domain with a 47-residue insertion - in the middle of the Peptidase_M20 split domain (Supplementary Table S2).

**Figure 1:**
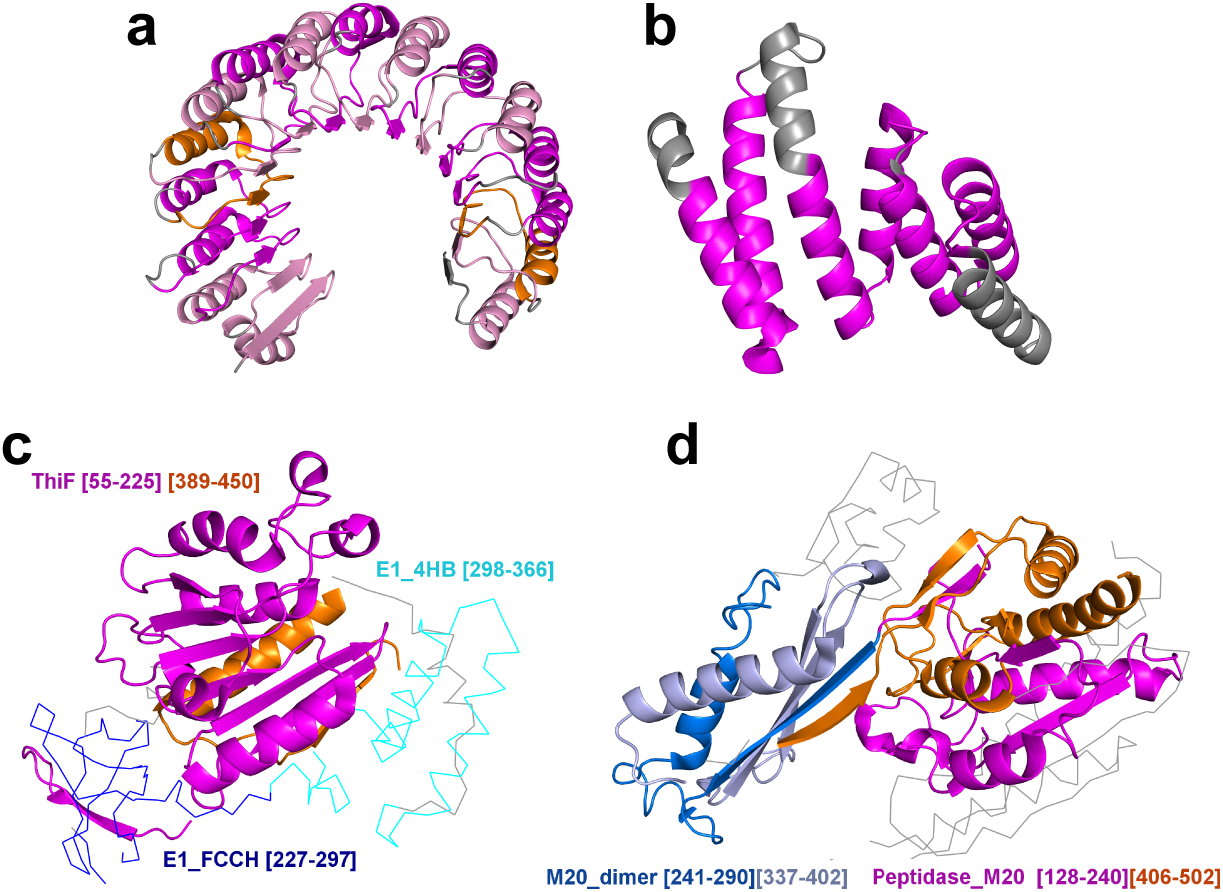
Assignment of Pfam domains to the human proteome. **a-b** Examples of repeat Pfam assignments in RINI (LRR repeats; PDB: 1Z7X) and AIP (TPR repeats; PDB: 4APO) proteins. Repeats assigned both in HuPfam and in Pfam are colored in orange, repeats assigned only in HuPfam are colored in magenta, and weak repeats in HuPfam are colored in pink. Regions with no Pfam assignments are colored in gray. **c-d** Examples of split Pfam assignments. ThiF split domain of UBA1 (PDB: 4P22) has two inserted domains: E1_FCCH (blue ribbon) and E1_4HB (cyan ribbon). CDNP1 has two split domains - Peptidase_M20 (magenta, orange) and M20_dimer (blue, light blue). Regions not assigned to Pfams are shown in gray ribbons.

We used the HuPfam assignments for all further analysis. Importantly, because Pfam domains are derived from a set of homologous proteins, their HMM profiles capture characteristics not only of human proteins but also of homologs from other species. If a PPI in the human network is not structure-supported directly through a complex structure of two interacting human proteins, it can still be structure-supported through an equivalent homologous complex from other species. Therefore, our Pfam domain-based approach makes the identification of structure-supported PPI in the network more robust to species-based sequence variation. We treat a human PPI as structure-supported even if the only evidence for the interaction are structures of homologous protein complexes from other species.

### The human structure-supported interactome is a subset of the total interactions

We used Pfam-based domain–domain interfaces to identify the structure-supported interactions in the human interactome. We constructed the human interactome by merging protein-protein interactions from three source interaction networks: i) the BioPlex network^2^ comprising more than 56,000 PPI generated by affinity purification followed by mass spectrometry (AP–MS), ii) a yeast two-hybrid network^1^ with over 14,000 binary PPIs, and iii) a curated literature-based network from Rolland *et al*.^1^ of about 11,000 high-quality binary interactions from seven PPI databases. The merged interaction network contains nearly 80,000 interactions among 13,600 proteins representing about two thirds of the encoded human proteome.

Using ProtCID, we first identified all the known structural domain–domain interfaces used by proteins in PPI by extracting Pfam-based^26^ interfaces in the biological assemblies of protein structures in the PDB (Supplementary Fig. S2). We assume a protein–protein interaction to be constituted by a set of one or more domain–domain interfaces, and for brevity refer to such PPI as domain–domain interactions. We defined a network PPI to be structure-supported if the interacting proteins contain one or more domains that enable them to interact through at least one known domain–domain interface (Fig. 2a). Thus, the structure-supported PPI are limited by the number of structurally resolved domain–domain interfaces. One-eighth of the PPI in the network are structure-supported (Fig. 2b). In contrast, only 3.25% of the interactions are structure-supported in a control PPI network generated by randomizing the nodes while preserving the edges found in the human interactome (Supplementary Fig. S3). As shown in Figure 2c, PPI found in multiple subnetworks are significantly enriched for structure-supported interactions (*P*≈0, hypergeometric test). To determine the effects of the size of interacting proteins on structural support for PPI, we computed the number of structure-supported interactions among interacting proteins of various domain lengths (Fig. 2d). Nearly half of the structure-supported interactions involve protein pairs containing two domains or lower on average. Even among interactions involving long multi-domain proteins (>10 average domains/interacting protein pair), about a quarter are structure-supported highlighting the advantage of using a modular domain-based approach to identify structural interfaces.

**Figure 2:**
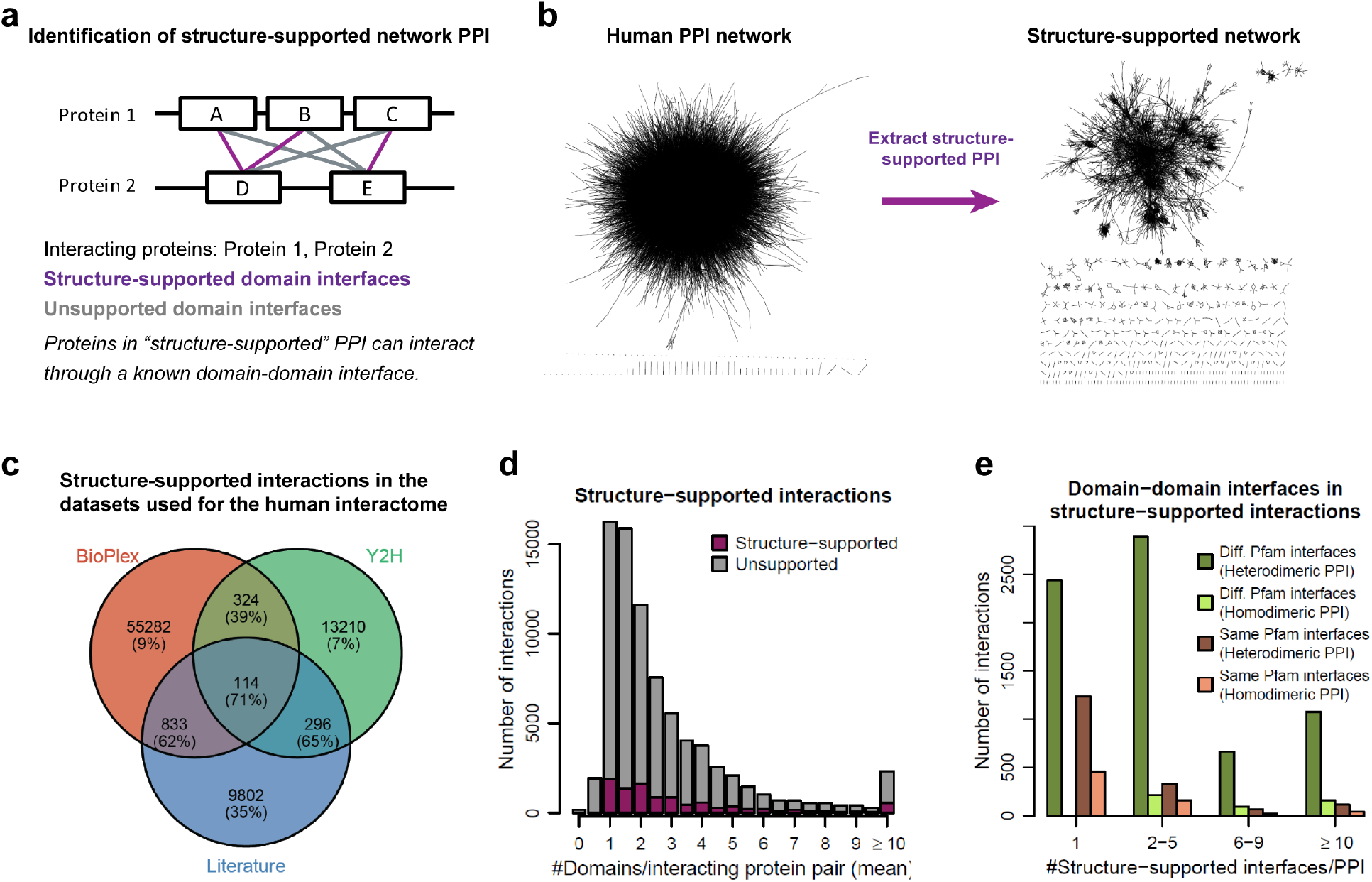
Characterization of the structure-supported PPI network. **a** Identification of structure-supported domain–domain interfaces in a PPI. **b** The structure-supported network is a subgraph of the human interactome and contains about 12.6% of the total PPI. **c** Distribution of PPI among the three interactome datasets used to construct the merged interaction network. Numbers in the parentheses represent the percentage of structure-supported PPI within each subset. **d** Number of structure-supported interactions among interacting proteins of various domain lengths. X-axis shows the average number of domains for every interacting protein pair. **e** Pfam domain–domain interfaces used by proteins in the structure-supported homodimeric and heterodimeric PPI in the network.

To evaluate the type of Pfam domain–domain interfaces used by proteins in PPI, we checked how often PPI in the network solely form interfaces between homologous domains (same Pfam) that are present in both interacting proteins (same-Pfam interfaces). Although same-Pfam interfaces are homodimers at the domain-level, they can be observed in 3D structures of both homodimeric and heterodimeric protein– protein complexes as different interacting proteins can have the same homologous Pfam (Supplementary Fig. S4). A quarter of all the structure-supported PPI are predicted to interact through same-Pfam interfaces (Fig. 2e). As expected, same-Pfam interfaces are more prevalent in homodimeric PPI (59.5%) compared to heterodimeric PPI (19.8%), both of which include multi-domain proteins. Remarkably, homodimeric interactions between multi-domain proteins predicted to form just one structure-supported domain–domain interface are all based on same-Pfam interfaces, indicating that protein homodimers prefer to interact through their identical domains when that is the sole interface forming the PPI. Moreover, nearly three quarters of the structure-supported heterodimeric PPI are based on domain– domain interfaces that are exclusively observed in heterodimeric protein complexes and are hence likely to be found in PPI involving different proteins.

Domain-domain interfaces in the ProtCID database come from both inter-chain interfaces and intra-chain interfaces in multi-domain proteins. Over 93% of the network PPI are structure-supported because the interacting proteins contain domains that are found in interchain domain–domain interfaces. A small fraction of the PPI (5.8%) are structure-supported only through intrachain domain–domain interfaces. The few remaining structure-supported PPI (1%) are based only on full-length chains from the PDB, *i.e*., structural interfaces between specific entire protein chains in the PDB. This occurs regions of a protein that interact with another protein are not covered by any Pfam assignment.

### The structure-supported network is scale-free and hierarchical like other biological networks

We calculated topological properties of the structure-supported network to characterize its overall organization. First, we evaluated the average degree of the nodes in the network. Degree of a node is the number of its direct connections in the network and hence represents the number of interaction partners for each protein. The average degree for the nodes in the structure-supported network is 3.35. The average degree of the Y2H structure-supported subnetwork (2.27) is lower compared to the BioPlex (2.70) and Literature (2.78) structure-supported subnetworks, likely because the Y2H interaction network was designed to be a homogenous network covering a large portion of the previously unexplored proteome. We also calculated the degree distribution of the nodes across the network. As shown in Supplementary Fig. S3a, degree distribution (*P*(*k*)) of the nodes in the structure-supported network obeys the power law, *P*(*k*)~*k*^−2.48^, where *k* is the node degree, illustrating their scale-free topology.^28^ Second, we calculated the transitivity for each node in the network, defined as the fraction of interconnectivity between all the nodes adjacent to each specific node. Transitivity of the nodes gradually drops with an increase in the number of protein interactions indicating a hierarchical network organization comprising highly connected proteins (or “hubs”) interacting with sparsely connected proteins (Supplementary Fig. S3b). However, some highly connected nodes (>15 direct interactors) demonstrate high transitivity suggesting the presence of dense subnetworks of interacting hub proteins. Third, we measured the fraction of the nodes in the Largest Connected Component (LCC), representing the biggest subgraph of nodes connected to each other by paths in the network. LCC fraction for the structure-supported network is only 0.63 highlighting the sparse structural data beyond the well-explored regions of the interactome.

We next identified all interactome PPI that are found as complexes in PDB structures (e.g., a co-crystal structure of a PPI in the network). These interactions constitute a smaller subnetwork comprising only 2% of the human interactome, or a subset (15%) of structure-supported edges based on all Pfam domain– based interfaces (Table 1). The literature PPI subnetwork has the highest fraction of co-crystallized PPI as it is derived from well-studied proteins that have a higher chance of being targeted for structural studies. If co-crystal structures of all human protein complexes were to be excluded when building the structure-supported interactome, the number of structure-supported interactions would only drop to 10.5% (compared to 12.6% when using all structures), highlighting the impact of homologous interfaces used for capturing the majority of PPI across species and paralogous genes (Supplementary Table S3). For all further analysis, we used the network of structure-supported interactions identified using Pfam domain-based interfaces, henceforth referred to simply as the structure-supported network.

**Table 1:** The structure-supported interaction network. Network characteristics of the structure-supported BioPlex, Y2H, Literature, and the overall merged networks. Numbers in the parentheses represent the corresponding values for the original networks and include PPI that are not structure-supported.

|  | **BioPlex +**<br>**Structure** | **Y2H +**<br>**Structure** | **Literature +**<br>**Structure** | **Merged +**<br>**Structure** |
| --- | --- | --- | --- | --- |
| Pfam-supported edges | 9.8% | 9.3% | 37.9% | 12.6% |
| Cocrystallized edges | 0.6% | 1.8% | 11.5% | 1.9% |
| Number of nodes | 4122 (10961) | 1146 (4303) | 3016 (5541) | 5996 (13591) |
| Number of edges | 5563 (56553) | 1298 (13944) | 4189 (11045) | 10055 (79861) |
| Average degree | 2.70 (10.32) | 2.27 (6.48) | 2.78 (3.99) | 3.35 (11.75) |
| LCC fraction | 0.54 (0.99) | 0.30 (0.95) | 0.50 (0.89) | 0.63 (0.99) |

### Structural interfaces highlight high-confidence and direct protein interactions

The structure-supported network is a subgraph of the human interactome and is constructed from evidence of existing homologous domain–domain interfaces for interacting protein pairs in the network. The structure-supported network is highly modular (modularity: 0.74), with densely interconnected protein subgraphs or “clusters” that have sparse connections to proteins in the other clusters. We used the Markov Cluster Algorithm (MCL) to divide the structure-supported network into smaller subgraphs of functionally related protein complexes (see Methods), or functional neighborhoods. Besides containing fewer proteins (nodes), proteins in the structure-supported network have fewer direct interactions (edges) on average (1.68 vs. 5.88 in the original network). We calculated the enriched GO biological process and molecular function terms for the highly-connected “hub” proteins with more than 15 direct interactions (Supplementary Tables S4-6) in the structure-supported network. The hub proteins are generally involved in the innate immune response (*P* = 1.4×10^-20^, hypergeometric test), regulation of mitotic cell cycle (*P* = 1.2×10^-16^), EGFR signaling (*P* = 3×10^-16^), and are enriched for protein binding (*P* = 1.1×10^-29^), ATP binding (*P* = 4.9×10^-14^), and kinase activity (*P* = 3.8×10^-13^). Growth factor receptor-bound protein-2 (Grb2), a key adaptor protein known to bind epidermal growth factor receptor (EGFR) leading to activation of RAS and its downstream kinases,^31^ has 61 interaction partners and is the most connected protein in the structure-supported network (Supplementary Table S4). Overall, only 11.5% of the proteins in the structure-supported network retain all the direct interactions from the original interaction network, but the structure-supported network edges represent high-confidence binary PPI. The discussion of the exosome complex below further illustrates this.

Figure 3a shows an example protein interaction cluster containing the human exosome, a multi-subunit 3’-5’ exoribonuclease complex involved in RNA processing and degradation.^32^ The exosome complex contains nine core subunits encoded by *EXOSC1-EXOSC9*, and commonly associates with other obligate proteins including the RNases Dis3L, PM/Scl-100 (*EXOSC10*), and the RNA binding protein C1D for processing of certain substrates. The human interactome contains 49 inter-protein interactions between the 12 proteins in the cluster compared to 31 interactions in the structure-supported network. In the human interactome, C1D interacts with ten subunits of the exosome complex, whereas the structure-supported interactome contains only one interaction. This is not surprising as all the C1D interactions are derived from the AP-MS experiments that report both indirect and direct interactions. The single edge in the structure-supported network, however, represents a direct physical interaction between PM/Scl-100 and C1D, an association that is experimentally validated through its role in pre-rRNA processing.^33^ While the interactions between the nine core subunits (blue edges) are captured in the reported cocrystal structure (PDB ID: 2NN6) of the human exosome complex, the interaction edges involving the three obligate proteins are based on domain-domain interfaces found in the homologous 12-subunit yeast exosome complex^34^ (PDB ID: 5C0W). Figure 3a show the structures of the yeast exosome complex and the nine core subunits (red) from the human exosome complex (C_α_-RMSD between yeast and human core subunits is 2.7 Å). The yeast subunits homologous to the human proteins PM/Scl-100, C1D, and Dis3L are colored in blue, yellow, and cyan, respectively.

**Figure 3:**
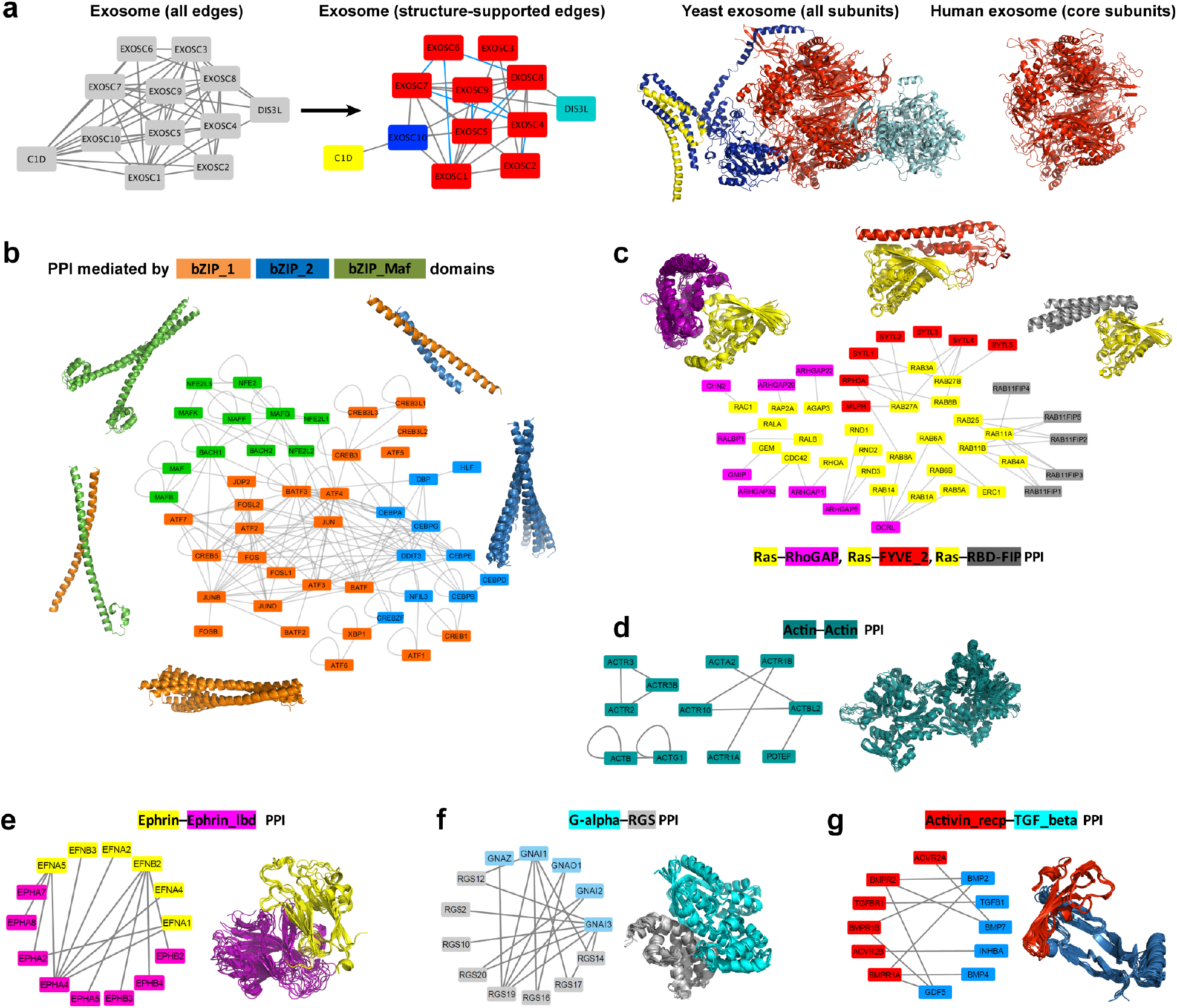
Extraction of PPI details from the structure-supported network. **a** Interaction of the pro-apoptotic protein C1D with the human exosome complex comprising ten subunits (EXOSC1-EXOSC10 and DIS3L). The local network of interactions in the original BioPlex network (left) are compared to the structure-supported network (right). Gray edges (Pfam-supported) in the structure-supported network are based on domain–domain interfaces, while the blue edges (cocrystallized) connect proteins known to interact in the cocrystal structure of the human complex. Structures of the 12-subunit yeast exosome complex (PDB ID: 5C0W) and the 9-subunit human exosome core complex (PDB ID: 2NN6) are also shown. **b-g** Examples of network PPI based on conserved domain–domain interfaces. Proteins are colored based on the domains they contribute to the domain–domain interfaces formed by the PPI.

To test the number of direct protein–protein contacts identified by the integration of structural interfaces in the human interactome, instead we calculated the number of PPI in the human interactome that are eliminated between proteins in structure-supported clusters. Use of Pfam domain-based interfaces to identify structure-supported interactions helps eliminate indirect interactions by reducing the number of edges (compared to the human interactome) in nearly 45% of the clusters with over ten interacting proteins. The average fraction of Pfam-based structure-supported PPI is similar across various protein interaction cluster sizes (0.94–0.98) and can even be as low as 0.5 in some clusters (Supplementary Fig. S6a, gray line and envelope, respectively). In contrast, the fraction of PPI involving direct co-crystallized protein pairs gradually declines with increasing protein interaction cluster size and drops from 0.17 in protein interaction clusters containing up to five proteins to 0.06 in clusters comprising 25-30 proteins, thus, highlighting the increased complexity for crystallization studies if only human proteins are used. Moreover, over half of the three-protein or larger clusters do not have even a single PPI between a cocrystallized human protein pair and hence can only be structurally modeled using domain-based interfaces from homologous proteins.

### Structural domain–domain interfaces offer insights into domains facilitating PPI

Nearly 40% of the PPI in the structure-supported network are based on single domain–domain interfaces within each protein involved in the PPI. For the remaining PPI, between protein pairs that interact through multiple domain–domain interfaces, structural data highlight the interacting domains. The top 5% most frequently used domain–domain interfaces cover over half of all structure-supported interactions (Supplementary Fig. S6b) and highlight the presence of widely reused interaction modes across PPI in the network. On the other hand, there are 6465 known structural interfaces involving the 5501 unique domains found in the human interactome proteins, but only 49.6% of those interfaces are used in at least one PPI in the network, suggesting significant gaps in the model human interactome.

Identification of similar domain–domain interfaces in PPI reveals conserved interaction motifs used by proteins in the structure-supported interactome. For example, Figure 3b shows the interaction subnetwork comprising 171 interactions between 46 basic region leucine zipper (bZIP) transcription factors, a family of proteins previously studied for their PPI specificities.^35,36^ The bZIP transcription factors in the subnetwork each contain one of the three bZIP Pfam domains – bZIP_1 (orange), bZIP_2 (blue), or bZIP_Maf (green). Remarkably, all the interactions between these proteins are localized to their bZIP domains and are covered by just five domain–domain interfaces. Only 15% of the PPI in the bZIP subnetwork are between protein homodimers, while over 70% of the bZIP subnetwork PPI are based on homodimeric Pfam domain–domain interfaces. Similarly, Figure 3c shows the structure-supported PPI network around proteins from the guanine nucleotide binding Ras superfamily – primarily Rab and Rho GTPases that regulate intracellular membrane traffic^37^ and actin dynamics.^38^ The Ras superfamily proteins (yellow) all contain Ras Pfam domains. The subnetwork captures their interactions with proteins containing (i) RhoGAP domains (purple), most of which inactivate the Ras proteins by enhancing their intrinsic GTPase activity,^39^ (ii) FYVE_2 domains (red), most of which are Rab effectors that regulate exocytosis,^40^ and (iii) RBD-FIP domains (gray), which modulate Rab11-dependent vesicle recycling.^41^ Despite the presence of several other domains in proteins within each group, all 47 interactions can be mapped to interfaces between the highlighted domains and Ras domains in their partner proteins.

Additional analysis reveals conserved domain–domain interface motifs used for PPI between proteins within each of the different functional classes and neighborhoods in the network. A few examples include PPI mediated by (i) Actin–Actin interfaces between actin and actin-related proteins (turquoise), primarily involved in cell motility, cytoskeleton maintenance, and gene regulation (Fig. 3d);^42^ (ii) Ephrin–Ephrin_lbd interfaces between ephrins (yellow) and Eph receptors (purple), which regulate several developmental processes (Fig. 3e);^43^ (iii) G-alpha–RGS interfaces between G protein α subunits (cyan) and RGS proteins (gray), which inactivate G proteins leading to rapid turnoff of GPCR signaling pathways (Fig. 3f);^44^ (iv) Activin_recp–TGF_beta interfaces between transforming growth factor (TGF) beta superfamily proteins (blue) and their receptors (red), which drive a wide range of cellular processes including growth, differentiation, and homeostasis (Fig. 3g);^45^ and (v) UQ_con–zf-C3HC4_2, UQ_con–zf-C3HC4_3, UQ_con– zf-C3HC4_4, UQ_con–zf-RING_2 interfaces between E2 ubiquitin-conjugating enzymes and E3 ubiquitin ligases, which are responsible for protein ubiquitylation (Supplementary Fig. S7).^46^ The structures of domain–domain interfaces can thus be used to identify which individual domains within the proteins participate in PPIs, an important step to understand the biophysical effects of the disease-associated variants that map to these proteins.

### The structure-supported network enables more detailed study of disease-enriched neighborhoods

Studies to understand the molecular basis of human diseases would benefit from the human structure-supported PPI network. Disease-causing mutations can occur in different regions of the protein that might change its structure and/or function, but it is challenging to decipher accurately how they affect function. The disease-causing effect may be direct if it occurs in the protein-protein interface, or indirect if it destabilizes the folded structure of an interacting domain, or the folding dynamics, expression pathways, or expression levels. To probe the structural effects of mutations on PPI, we mapped 38,264 literature-reported disease-causing mutations^47^ that fall within the protein-coding regions of proteins in the network (see Methods). Nearly 72% of these mutations map to Pfam domains within proteins (nodes) in the human interactome, and the remaining fall in protein regions outside domains (Fig. 4a, top). This number rises to over 83% when considering both point mutations within domains and insertions/deletions that terminate or shift the reading frame, thus affecting the downstream domains. We next evaluated the effects of the mutation-disrupted domains in proteins on PPI (edges) in the human interactome. Over one third of the mutations occur in domains that participate in one or more structure-supported interactions in the human interactome and hence could affect those PPI (Fig. 4b-c). 12% of the mutations are predicted to not directly affect interactions because they fall within protein regions that are not involved in any interfaces in structure-supported PPI, and the remaining 55% of mutations map to PPI that are not structure-supported (Fig. 4a, bottom). In the following examples, we discuss examples of disease-causing mutations that affect structure-supported interactions.

**Figure 4:**
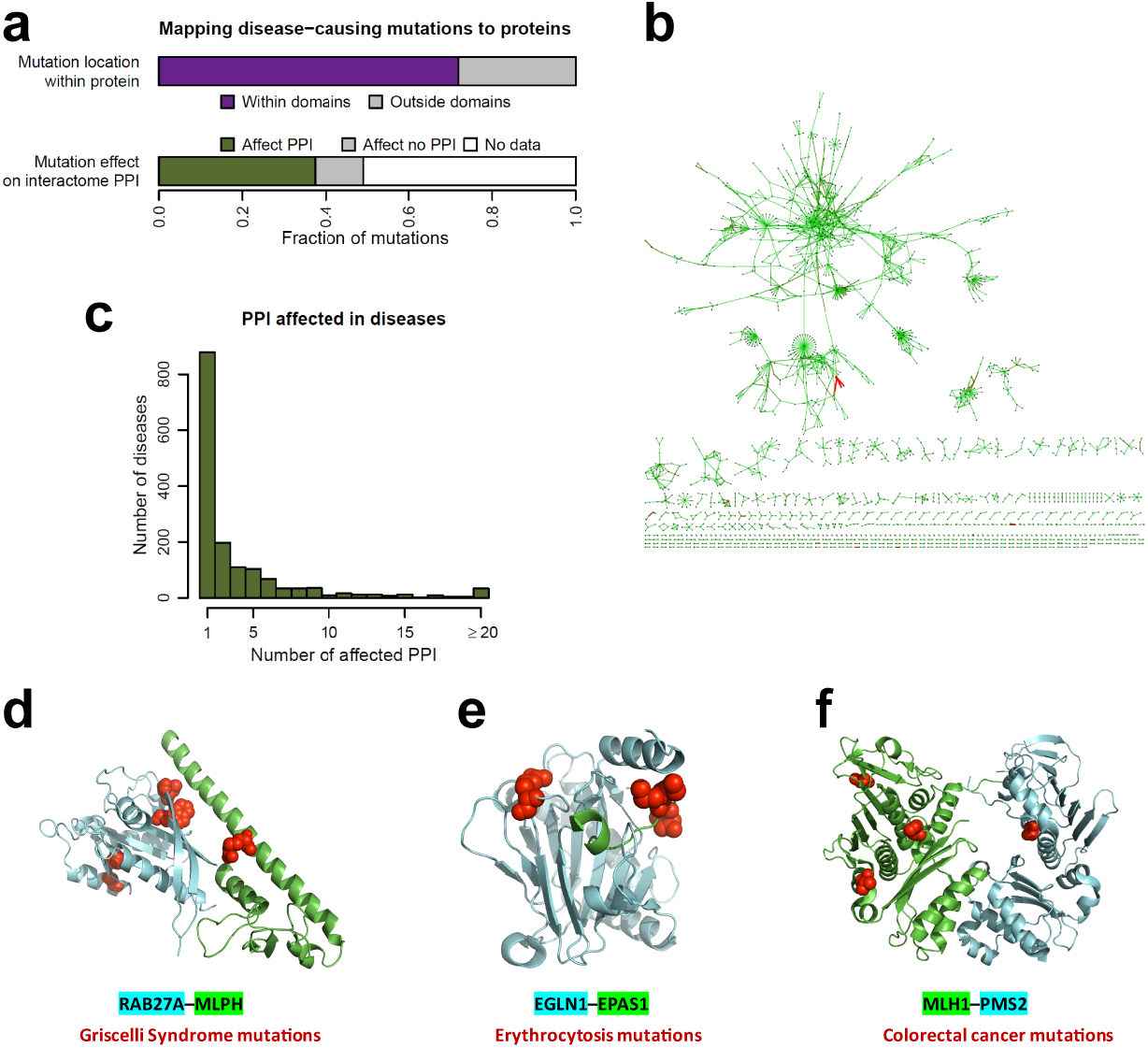
Estimation of the effects of disease-causing mutations. **a** Fraction of the known disease-causing mutations within the protein-coding regions that (i) fall within or outside domains, and (ii) are predicted to affect structure-supported PPI in the interactome. **b** Network of the interactome PPI predicted to be affected by at least one disease-causing mutation. **c** Number of diseases with known mutations predicted to affect PPI in the interactome. **d-f** Examples of PPI where disease-causing mutations (red spheres) map to domain–domain interfaces predicted to mediate the interaction.

Figure 4d shows a model of the interaction between Rab27A (*RAB27A*) (cyan) and Slac2-a (*MLPH*) (green) known to be involved in docking of multi-vesicular endosomes at the plasma membrane in the exosome secretion pathway.^48^ While Rab27A contains one Ras domain, Slac2-a contains one FYVE_2 domain and two Rab_eff_C domains. Structural support for the Rab27A–Slac2-a interaction is based on the conserved Ras–FYVE_2 domain–domain interface observed in the structures of the rat Rab3A–Rabphilin 3A (PDB ID: 1ZBD), mouse Rab27B–Slac2-a (PDB ID: 2ZET), and mouse Rab27A–human Slp2 (PDB ID: 3BC1) complexes. Structural domain–domain interfaces thus localize the PPI to the Ras–FYVE_2 interface, a finding supported by existing mouse immunocytochemical and cell-based studies indicating the involvement of the two domains in binding. Rab27A I44T, W73G missense mutations found in Griscelli syndrome type 2 patients,^50^ and Slac2-a R35W missense mutation found in Griscelli syndrome type 3 patients^51^ localize to the Ras–FYVE_2 domain interface (Supplementary Fig. S8) and have been shown to abrogate the Rab27A– Slac2-a interaction. Three other Griscelli syndrome-associated point mutations (A87P, L130P, A152P) map to the Ras domain in Rab27A but are not located at the Ras–FYVE_2 domain interface, making their effects difficult to predict; only one of these mutations (L130P) has been shown to affect the Slac2-a interaction.^52^ Finally, three known Griscelli syndrome-associated terminal mutations (at positions Q116, Q118, R184) are also predicted to disrupt the Ras domain and affect the Rab27A–Slac2-a interaction.

Figure 4e presents a model of the interaction between PHD2 (*EGLN1*) (cyan) and HIF2A (*EPAS1*) (green) leading to hydroxylation and eventual degradation of HIF2A, an important step in the maintenance of oxygen homeostasis.^53^ Although PHD2 contains two Pfam domains (zf-MYND, 2OG-FeII_Oxy_3) and HIF2A five Pfam domains (HLH, PAS, PAS_11, HIF-1, HIF-1a_CTAD), the PPI is localized to the 2OG-FeII_Oxy_3– HIF-1 domain–domain interface previously observed in structures of the human PHD2–HIF1A complexes (PDB IDs: 5L9B, 3HQR, 3HQU).^54^ Four erythrocytosis-associated mutations (M535V/T, G537W/R) map to the HIF-1 domain in HIF2A and are predicted to affect the PPI as they are located at the 2OG-FeII_Oxy_3– HIF-1 domain–domain interface, an observation supported by functional characterization of the disease-associated variants at these mutation sites.^55^ One erythrocytosis-associated point mutation in PHD2 (R371H), which maps to the 2OG-FeII_Oxy_3 domain is proximal to the 2OG-FeII_Oxy_3–HIF-1 interface of the PHD2–HIF2A interaction and has already been shown to impair the PPI.^56^ Additionally, one known erythrocytosis-associated terminal mutation (at pos. Q377) is predicted to disrupt the 2OG-FeII_Oxy_3 domain and affect the PHD2–HIF2A interaction.

The DNA mismatch-repair (MMR) complex of MutL protein homolog 1 and the mismatch repair endonuclease PMS2 (MLH1-PMS2) provides an illustrative example of how matching homologous proteins from different species can elucidate protein-protein interactions and disease associations of human proteins (Fig. 4f). Constitutional mutations in MMR-related proteins cause hereditary nonpolyposis colorectal cancers (HNPCCs, or Lynch syndrome), and are associated with substantially increased risks of colorectal and endometrial cancers, and with increased risk of multiple other cancers.^57^ Mutated MMR-related genes rank among the most widely recognized as predictive for hereditary cancers.^58^ MMR proteins form homodimeric complexes in bacteria while in eukaryotes they are heterodimers that are homologous to the bacterial homodimers. The bacterial homodimeric mutator S (MutS) complexes detect DNA base-base mismatches (MutSα) or loops caused by insertions/deletions (MutSβ) and initiate the remaining MMR reactions at the site. In a second step, the bacterial MutLα is recruited to MutS in an ATP-dependent manner, where it generates nicks into the DNA and engages DNA excision and replicative repair systems downstream. Eukaryotic homologs are the MSH2-MSH6 heterodimer for MutSα, and the MSH-2-MSH3 heterodimer for MutSβ. MutLα homologs are the heterodimers Mlh1-Pms1 in yeast and MLH1-PMS2 in humans.^59^ Consequently, mutations in MLH1, MSH2, MSH6, and PMS2 are the cause for or associated with many cancers. An important goal is to classify the pathogenicity of familial mutations and variants of uncertain significance (VUS) in MMR proteins to inform treatment approaches for carriers.^60^ Classification systems for such variants have been proposed^57^ and are an active area of research.

Several groups have constructed homology models of MMR protein structures to support the assessment of novel MMR protein VUS. MLH1 and PMS2 heterodimerize in two domains that are linked by unstructured linkers in each protein chain, respectively. The heterodimer formed by the N-terminal domains provides ATPase activity whereas the C-terminal domains are necessary for heterodimerization and endonuclease activity.^61^ To aid assessment of 15 VUS (5 each for MSH2, MLH1, and PMS2), we previously modeled missing loops in the structure of the human MSH2/MSH6 heterodimer.^62^ Depending on where a mutation is located within the protein complex, we reported that complementary computational and experimental assessments can help clarify the InSiGHT class of a VUS from class 3 (uncertain pathogenicity) to class 2 (likely not pathogenic) or 4 (likely pathogenic). Similarly, Köger, Plotz et al. assessed nine MLH1 VUS through in vitro biochemical and functional assays as well as structural analysis of homology models of the MLH1/PMS2 N-and C-terminal heterodimers.^63^ The authors conclude that in the absence of sufficient clinical data, which is viewed as the strongest basis for pathogenicity assessment,^60^ their combination of in vitro assays and structure analysis is a suitable alternative to assess the pathogenicity of VUS. In summary, interfaces within the structure-supported PPI network can help elucidate the effects of disease-associated mutations on PPI.

### Peptide-binding domains help predict PPI involving intrinsically disordered regions

In the previous sections, we identified structure-supported PPI in the human interactome based on the presence of Pfam domains known to interact directly via domain-domain interfaces as shown by structures in the PDB. However, a majority of protein residues in the human proteome do not form structural domains but are enriched in polypeptide regions without a defined three-dimensional structure. These regions are commonly referred to as intrinsically disordered regions (IDRs).^64^ To estimate the prevalence of IDRs in proteins in the human interactome, we used the MobiDB^65^ database to obtain the list of disordered residues in proteins of the network. Over 83% of the human interactome proteins contain at least 20 disordered residues each, and nearly 15% of the proteins contain over 300 disordered residues. We also evaluated the average number of disordered residues in the interacting proteins for each PPI in the human interactome (Fig. 5a). Nearly 95% of the PPI comprise protein pairs containing over 20 disordered residues per protein on average (all bins above 20 in Fig. 5a). Proteins that are mostly disordered (> 50% disordered residues) barely interact with each other (3% of the overall interactions). Therefore, a substantial proportion of the PPI that involve IDRs are likely to be between a structured domain from one protein and an IDR from the other.^66^ Complex structures of such interactions are typically solved with peptide fragments of the longer IDR-containing protein. In other cases, the structured domain really does bind a peptide. We define these PPI to be represented by domain–peptide interfaces, for brevity referred to as domain–peptide interactions.

**Figure 5:**
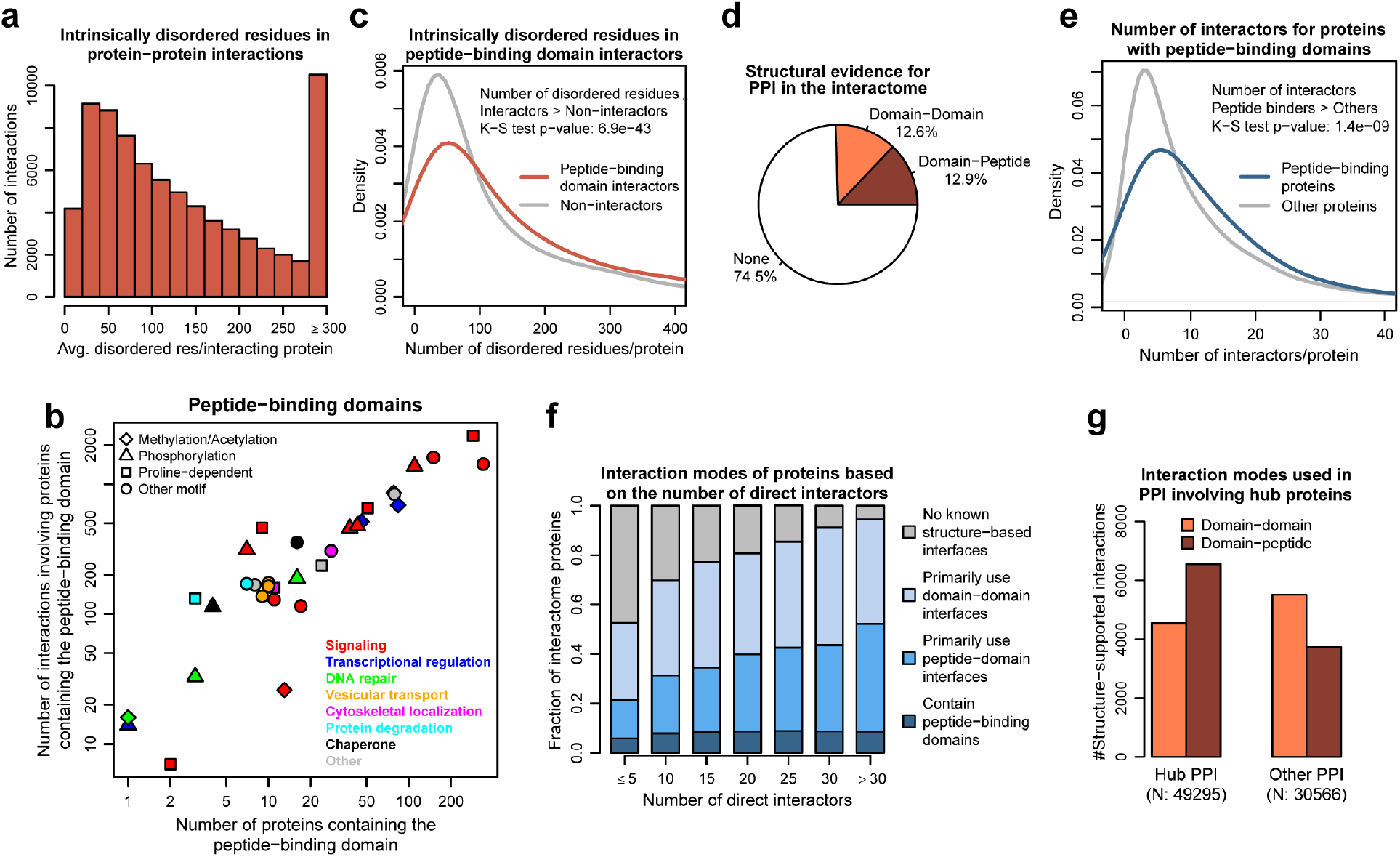
Identification of PPI mediated by peptide-binding domains. **a** Distribution of the number of PPI (edges) in the network connecting proteins (nodes) with different numbers of intrinsically disordered residues. The number of disordered residues involved in each PPI is calculated by averaging the disordered residues in its two interacting proteins. **b** Peptide-binding domains in the human interactome. Shapes represent the peptide recognition motifs of the domains and colors highlight the primary functions of the proteins containing the domains. **c** Kernel density estimates of the number of disordered residues in proteins that interact with peptide-binding proteins (interactors) and the remaining interactome proteins (non-interactors). **d** Percentage of structure-supported PPI based on domain–domain and domain–peptide interfaces. **e** Kernel density estimates of the number of direct interactors for proteins with and without peptide-binding domains. **f** Structural interaction modes used by the proteins in the network depending on the number of direct interactors. **g** Structural interaction modes used in PPI involving hub (>30 direct interactors) and other proteins.

Identifying structure-supported PPI based on domain-domain interactions and domain–peptide interactions within the human interactome are conceptually different. For the former, both protein nodes have Pfam domains derived from homologous sequences, and the connecting edge is defined by a ProtCID-classified biological interface between those domains. For the latter, only one node has a Pfam domain, and the connecting edge is defined by the peptide interaction preferences of that Pfam domain based on ProtCID domain–peptide clusters. The second node contributing the “peptide” cannot be reliably assigned any Pfam domain (or any other domain or motif) based on homology because of the short sequence. Because of this conceptual difference, our superposition of Pfam domain–domain interactions with the model human interactome allows for a well-defined count of the number of edges and nodes in the structure-supported network, but the number of structure-supported domain–peptide interactions can only be estimated.

To estimate the PPI structure-supported through potential domain–peptide interfaces in the human interactome, we first identified the commonly used peptide-binding Pfam domains in human proteins. We classified peptide-binding domains into two categories: i) enzymatic domains that catalyze modification of protein residues upon binding, and ii) non-enzymatic domains that do not modify protein residues upon binding. Enzyme–substrate interactions are mostly transient and are hence difficult to capture using Y2H or AP-MS methods commonly used for large-scale detection of PPIs.^67^ To estimate the prevalence of enzyme–substrate interactions in the human interactome used for this study, we accumulated the dataset of all enzyme–substrate interactions based on residue modifications reported in proteins in the UniProt database^27^ (Supplementary Table S7). Nearly 89% of the 2368 enzyme–substrate interactions in this dataset involve pairs of proteins (nodes) that are both part of human interactome, but only 9.2% of the interactions (edges) between them are captured in the human interactome. Moreover, even among the enzyme–substrate edges that are captured in the human interactome, over 60% are between proteins containing domains that would enable them to interact through a domain–domain interface. Since enzyme–substrate interactions are under-represented in the model human interactome used for the study, we excluded them and focused on interactions involving non-enzymatic peptide-binding domains for the remaining analysis.

To identify the commonly used non-enzymatic peptide-binding domains in the human interactome, we first extracted all domain-peptide interfaces in the PDB and clustered structurally similar domain–peptide interfaces for each Pfam domain (see Methods). After filtering out the enzymatic domains, we inspected the domain–peptide clusters of domains involved in more than 100 interactions in the network. We classified a Pfam domain as “peptide-binding” if peptide recognition/binding is an important component of its primary function. We also retrieved some less frequent peptide-binding domains in the interactome if they belonged to the same Pfam clans^26^ as the commonly used peptide-binding domains. We identified 52 peptide-binding domains primarily involved in signaling (23 domains), transcriptional regulation (12), DNA repair (3), vesicular transport (3), protein degradation (2), chaperone activity (2), and cytoskeleton-associated localization (2) (Table 2). Domains of the EF_hand clan, found in 339 proteins, are the most common peptide-binding domains in the proteome (nodes), while domains of the SH3 clan that participate in over 2350 interactions are the most widely used peptide-binding domains in the interactome (edges, Fig. 5b).

**Table 2:**
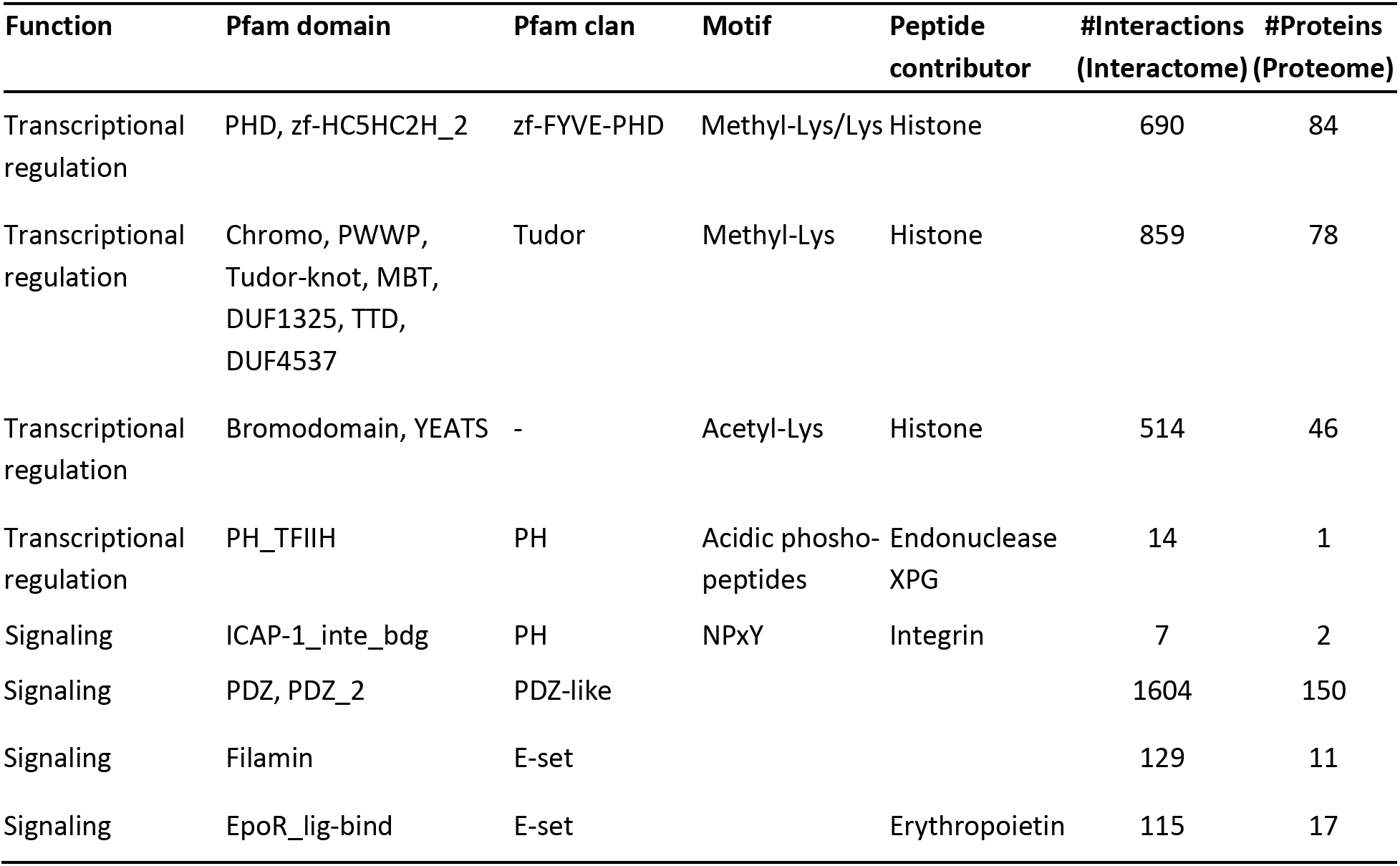
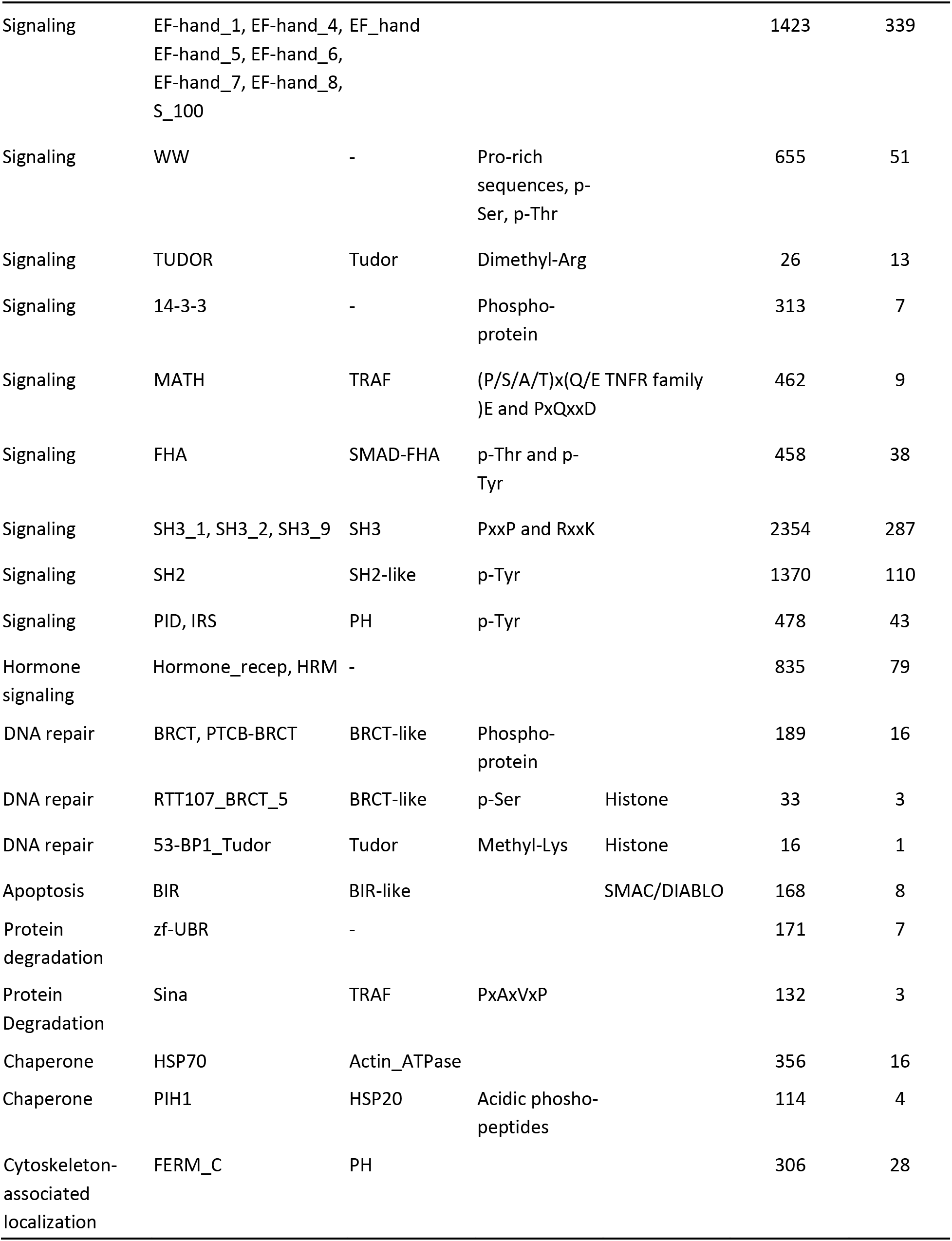
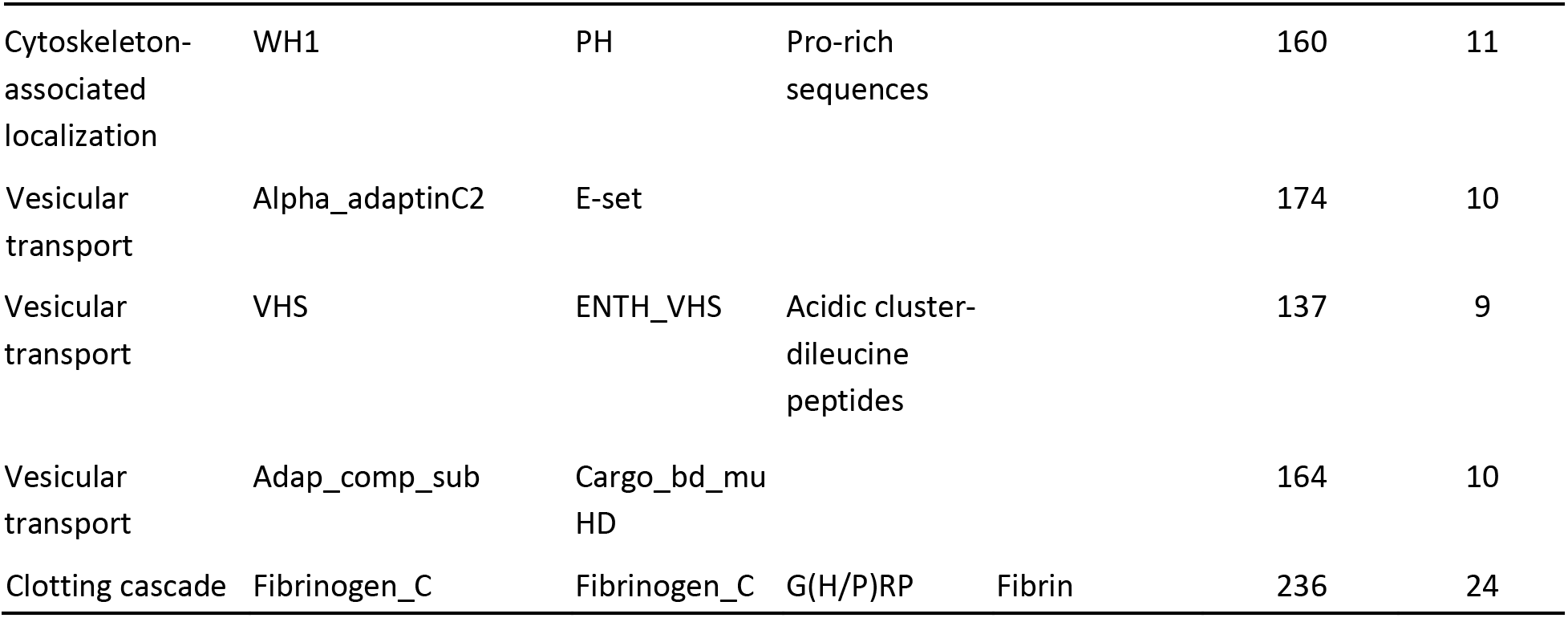
Peptide-binding domains in the human interactome. List of the identified structural peptide-binding domains grouped based on their recognition motifs and the primary function of the proteins containing the domains. Some domains are also specific to proteins contributing the peptide regions. The number of interactions involving proteins containing the peptide-binding domains in the interactome, and the number of proteins with the peptide-binding domains in the proteome are also listed.

We divided the peptide-binding domains into three broad categories based on their peptide recognition motifs: (i) domains that recognize post-translationally modified (PTM) sequences, which contain amino acids such as methyl-Lys, acetyl-Lys, phospho-Ser etc., (ii) proline-dependent domains that bind proline-containing sequences, and (iii) domains that bind other motifs. Domains of the SH2-like clan, which primarily bind regions that contain phospho-Tyr are the most widely used PTM-specific peptide-binding domains (1370 interactions), domains of the SH3-like clan are the most prevalent proline-dependent (2354 interactions) domains, and domains of the PDZ-like clan are the largest among the other peptide-binding domains (1604 interactions). Nearly 44% (23/52) of the peptide-binding domains specifically bind PTM sequences, contributing over 37% (4559/12231) of all PPI that involve domain–peptide interfaces (Supplementary Table S8).

Next, we characterized the peptide side of the domain–peptide interactions in the human interactome by comparing the prevalence of IDRs between proteins that do interact with peptide-binding domain proteins and those that do not. Figure 5c compares the distributions of the number of disordered residues in proteins (interactors), which interact with peptide-binding domain proteins with the remaining proteins in the human interactome (non-interactors). As expected, interactors of peptide-binding domaincontaining proteins have significantly more disordered residues compared to the non-interactor proteins across the human interactome (184 vs. 143 residues on average) (*P* = 6.9×10^-43^, Kolmogorov-Smirnov test).

### Domain–peptide interactions are as frequent as domain–domain interactions across the human interactome, but they dominate PPI involving hub proteins

In addition to 12.5% of the human interactome PPI that are structure-supported based on domain– domain interfaces, in our estimate 12.9% of the PPI involve proteins that contain at least one peptide-binding domain and are therefore likely to interact through domain–peptide interfaces (Fig. 5d). As demonstrated in Figure 5e, proteins with peptide-binding domains have more interaction partners compared to proteins that do not contain such domains (14 vs. 12 direct interactors on average) (*P* = 1.4×10^-9^, Kolmogorov-Smirnov test). We next calculated the interaction modes used by the proteins in the network depending on the number of their direct interactors (Fig. 5f). We classified protein nodes into four groups depending on their interaction modes: (i) proteins with peptide-binding domains that can interact with peptide regions in their interaction partners (dark blue), (ii) proteins that primarily interact through domain–peptide interfaces by using the peptide-binding domains of their interaction partners (blue), (iii) proteins that primarily interact through domain–domain interfaces (light blue), and (iv) proteins involved in PPI that do not have any structure-supported interfaces (gray). The percentage of proteins with peptide-binding domains increases from 5.5% among proteins with five or fewer interactors to 8.5% in proteins with 10-15 interactors and remains the same with further increase in number of interacting partners. Proteins with fewer direct interactors primarily interact through domain–domain interfaces rather than domain–peptide interfaces. Remarkably, the percentage of IDR-containing proteins that primarily interact with peptide-binding domains of their partners gradually trends from 12.8% in proteins with five or fewer interactors to 42.8% in proteins with over 30 interactors.

Hub proteins (> 30 direct interactors) are enriched for proteins that primarily interact through the peptide-binding domains in their interaction partners (*P* = 6.2×10^-47^, hypergeometric test). Interactions involving hub proteins are significantly enriched for domain–peptide interfaces (*P* = 6×10^-6^, hypergeometric test), whereas the remaining PPI (up to 30 direct interactors) are dominated by domain–domain interfaces (*P* = 2.6×10^-283^, hypergeometric test) (Fig. 5g). Specifically, hub protein interactions are enriched for structure-supported interactions involving their IDRs with PTM-and proline-containing sequences (*P* = 1.5×10^-14^ and 6×10^-8^, respectively, hypergeometric tests) that are recognized by peptide-binding domains of their many interaction partners, but are not enriched for interactions with other peptide-binding domains (*P* = 0.99, hypergeometric test) (Supplementary Table S8).

### Structural insights into the histone and BRCA1 PPI structure-supported subnetworks

We illustrate the biological utility of domain–domain and domain–peptide interactions on two examples of structure-supported subnetworks, the histone interactome and various BRCA1–neighbor interactions. The structure-supported histone interactome is dominated by domain–peptide interactions involving PTM-containing regions in the histone tails. Figure 6a shows the structure-supported interaction subnetwork around histones (gray) that are core components of the nucleosome.^68^ The subnetwork contains 76 PPI between 57 proteins. While the 19 histones interact with each other through structure-supported domain–domain interfaces (gray edges), the remaining 38 proteins shown in the subnetwork each contain at least one histone tail-specific peptide-binding domain and are hence predicted to interact with the histone tails through domain–peptide interfaces (black edges).

**Figure 6:**
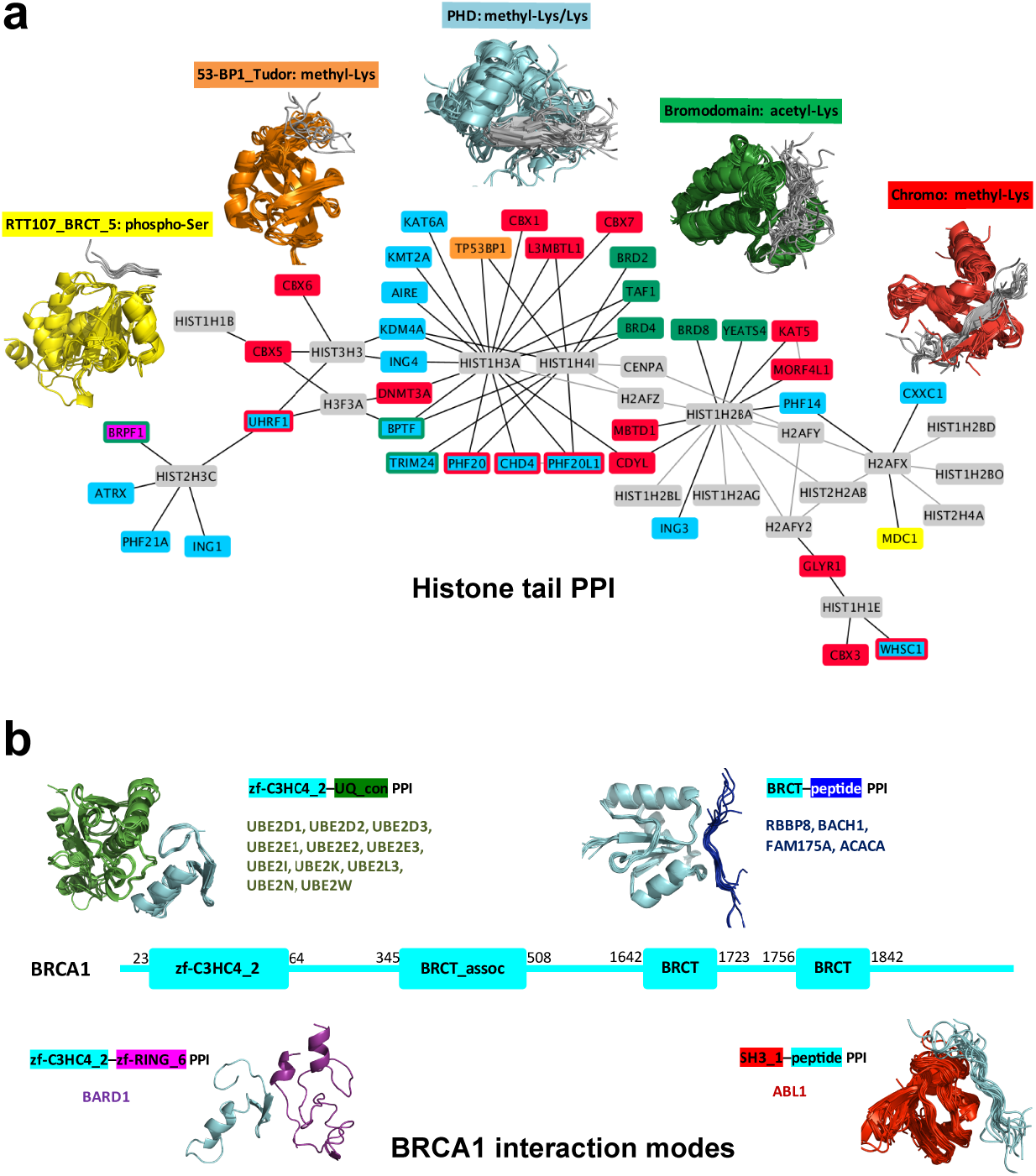
Interaction modes used by proteins in PPI. **a** Structure-supported interaction subnetwork between histones (gray) and proteins with histone tail-specific peptide-binding domains. Gray edges represent domain–domain interfaces and black edges represent domain–peptide interfaces. Nodes are colored based on the domains within each protein with colored borders indicating presence of more than one class of peptide-binding domains. The histone tail motifs recognized by the peptide-binding domains are also shown. **b** Examples of structure-supported BRCA1 (cyan) interactions mediated by various domain–domain and domain– peptide interfaces. BRCA1 Interactors are colored based on the domains or peptide regions they contribute to the PPI. Numbers represent the residue ranges for the four BRCA1 Pfam domains. The N-terminal zf-C3HC4_2 domain in BRCA1 forms domain– domain interfaces with proteins that support its E3 ubiquitin ligase activity. The C-terminal BRCT domains are peptide-binding domains and help BRCA1 bind phoshoproteins required for its role in tumor suppression and modulation of lipid synthesis. BRCA1 also interacts with other peptide-binding proteins through its peptide regions.

The histone tail-binding proteins can be broadly classified into five categories depending on the primary function and/or the epigenetic marks recognized by their peptide-binding domains. Most of the histone-specific peptide-binding domains are involved in transcriptional regulation, with PHD (cyan), Chromo (red), and Bromodomain (green) representing the classes of domains that recognize methyl-Lys/Lys, methyl-Lys, and acetyl-Lys in histone tails, respectively (see Table 2). The other two classes of histone-specific peptide-binding domains, 53-BP1_Tudor (orange) and RTT107_BRCT_5 (yellow), participate in DNA repair and recognize methylated-Lys and phospho-Ser in histone tails, respectively. While most of the histone tail binding-proteins (30/38) have either one of more copies of a single class of peptide-binding domain, eight of them have peptide-binding domains belonging to two or more classes and hence may recognize multiple histone modifications. Peregrin (*BRPF1*) is the only protein in the subnetwork with peptide-binding domains from three different classes and is a known chromatin regulator responsible for recognizing multiple epigenetic marks during embryonic development and cell proliferation.^69^ Once the proteins localize to the histone tails through their peptide-binding domains, the proteins can modify chromatin using their other enzymatic domains or recruit additional chromatin modifier proteins.^70^ For example, KAT5 and KAT6A both have MOZ_SAS acetyltransferase Pfam domains C-terminal of their histone tail-binding domains and are catalytic subunits of multi-component acetyltransferase complexes. Additionally, some histone-binding proteins in the subnetwork recognize modifications on multiple histones and can associate with other histone-binding proteins to aid chromatin compaction. For example, L3MBTL1 protein has three histone tail-binding MBT domains and can associate with the HP1 protein (*CBX*) to form a multivalent chromatin binder and help compact nucleosomal arrays.^71^ Therefore, systematic characterization of histone-specific peptide-binding domains helps identify proteins that form the core histone recognition modules of various transcriptional regulation and DNA repair complexes.

Figure 6b shows examples of structure-supported interactome PPI involving BRCA1 (cyan) that form various domain–domain and domain–peptide interfaces. BRCA1 has four Pfam domains – zf-C3HC4_2, BRCT_assoc, and two BRCT domains. The N-terminal zf-C3HC4_2 domain is predicted to be involved in PPI through domain–domain interfaces with (i) UQ_con domains in 11 different E2 ubiquitin-conjugating enzymes (green), and (ii) zf-RING_6 domain in the BRCA1-associated RING domain (BARD1) protein (purple). These PPI are known to play a central role in the E3 ubiquitin ligase activity of BRCA1.^72^ The C-terminal BRCT domains are peptide-binding domains and help BRCA1 interact with phosphorylated isoforms of repair proteins CtIP (*RBBP8*), FancJ (*BACH1*), and Abraxas (*FAM175A*) (blue) through domain– peptide interfaces. Phosphoprotein binding by the BRCT domains is known to be critical for BRCA1’s DNA damage response and tumor suppression.^73^ BRCA1 also modulates lipid synthesis through its interaction with phosphorylated acetyl-CoA carboxylase (*ACACA*) involving BRCT domains.^74^ Finally, BRCA1 also interacts with other proteins containing peptide-binding domains through its intrinsically disordered regions. For example, c-Abl (*ABL1*) (red) has an SH3 peptide-binding domain that directly interacts with the PXXP motif in the C terminus of BRCA1 impacting cellular response to DNA damage.^75^ Thus, integration of domain-domain and domain-peptide structural interfaces reveals the diverse interfaces used by BRCA1 to directly interact with its various partner proteins. The structure-supported interactome can thus be used to guide more detailed studies on the different interaction modes used by the proteins in PPI with their diverse interaction partners.

## Discussion

We describe modular integration of both structure-based domain–domain and domain–peptide interfaces with the human interactome. One-eighth of the human PPI are structure-supported as the interacting proteins contain domains known to form domain–domain interfaces in 3D structures of biological assemblies of proteins in the PDB. These biological interfaces represent high-confidence binary PPI and help eliminate indirect interactions from the human interactome in about half of the PPI clusters with ten or more interacting proteins. Use of domain-based structural interfaces also helps localize the PPI to domains within the interacting proteins as nearly 40% of the structure-supported PPI are based on single domain–domain interfaces, narrowing the PPI to one domain within each interacting protein. Almost three quarters of the literature-reported disease-causing mutations (HGMD database) in the protein-coding regions map to domains enabling more detailed analyses of the effects of mutations on structure-supported PPI. Finally, we identified the most common non-enzymatic human peptide-binding domains. Assuming that each peptide-binding domain binds to at least one IDR or peptide region, we estimate that the number of structure-supported domain-peptide interactions is comparable to that of the domain-domain interactions, and that inclusion of domain–peptide interactions approximately doubles the number of structure-supported interactions. Overall, structure-supported interactions offer a higher-resolution view of human PPI allowing prioritization of more in-depth biophysical or functional studies of proteins of interest and under consideration of their network neighbors.

The number of structure-supported PPI in the human interactome is limited because of two main reasons. First, the structure-supported network edges mostly represent direct physical contacts and ignore the indirect interactions between proteins in the human interactome. However, even structure-supported edges can sometimes overestimate the number of binary interactions, e.g., in multi-protein complexes composed of structurally similar subunits that all have the same homologous domains. Second, the structural coverage of the current domain-based interfaces in protein complexes in the PDB is not complete. Protein complexes are more challenging to crystallize, especially large complexes with multiple protein subunits. Also, in most cases only smaller segments (domains or IDRs) of the full-length proteins in the complex are crystallizable. With advancements in cryo-electron microscopy (cryo-EM) revolutionizing structural studies of large protein complexes,^76^ the diversity of known interfaces is expected to expand. This can boost structural coverage of PPI in the human interactome, especially if newly discovered domain–domain interfaces are widely used throughout the network.

Since the structure-supported network localizes the PPI to domains within the interacting proteins, it can be used to guide functional studies of disease-associated mutations that map to interfaces and disrupt PPI more frequently than mutations away from interfaces.^13,77^ Importantly, we provide links to clusters of homologous domain–domain interfaces observed not just within the protein biological assemblies but also within their other crystal forms and/or secondary interfaces to enable studies of mutation effects on other potential biologically-relevant interfaces. Even if the mutations do not locate to any of the interfaces, use of domain-based interfaces as the foundational blocks for PPI allows prioritization of experimental studies to domains containing the relevant mutations. We increased the robustness of our Pfam assignments to human protein sequences allowing us to use the cross-species homology inherent in Pfam domain definitions and expand the structure-supported coverage of the human proteome. Encouragingly, the minor drop in structure-supported human PPI when using domain-based interfaces exclusively from non-human protein complexes highlights the abundance of structural data from other species that can be exploited to design studies to understand the underlying biology and/or the effects of human disease-associated mutations.

We specially focused on PPI mediated by intrinsically disordered regions (IDRs) through identification of potential PPI involving peptide-binding domains. Our prediction represents an estimate of the number of domain–peptide mediated PPI in the human interactome and is comparable to the previous 15-40% estimate.^20^ Although we highlighted the peptide motifs targeted by the peptide-binding domains (where known), we did not predict the exact peptide regions involved in each domain–peptide mediated PPI as the consensus sequences are highly variable and are often dependent on post-translational modifications (PTMs). Hub proteins (nodes), especially those that interact through only one or two interfaces have been previously predicted to contain more IDRs than other proteins in the proteome.^78^ Our analyses further show that PPI (edges) involving hub proteins are also enriched for domain–peptide interactions, primarily involving the PTM and proline-containing IDRs in hub proteins and the peptide-binding domains in their interaction partners.

High-throughput interactome mapping studies based on biophysical assays typically include a statistical score-based filtering step to screen out false positive interactions observed in the individual experiments. Structural interfaces can be used to rescue potential biologically-relevant PPI that could be eliminated during such filtering. Furthermore, the modular structural domain-based approaches developed for this study can be quickly expanded to allow ortholog comparisons in interactomes of other model organisms.^79,80^ Our study presents a resource to generate hypotheses and drive design and prioritization of experimental studies of clinical significance.

## Materials and Methods

### Human proteome Pfam domain assignment

The procedure used to assign Pfams to human proteins was similar to the one described in our assignments of Pfams to proteins in the PDB (Supplementary Fig. S1).^25^ For each human protein, we used the Uniprot sequence, two consensus sequences, and one hidden Markov model (HMM) to find Pfam alignment hits. First, for each sequence, we ran PSI-BLAST on the UniRef90 database from UniProt (ftp://ftp.uniprot.org/pub/databases/uniprot/uniref/uniref90/). Two consensus sequences were derived from each PSI-BLAST^81^ profile, of which “percentage consensus sequence” is composed of the most frequent residues at each residue position, and “PSSM consensus sequence” is composed of highest scoring amino acid at each position. We also ran HHblits^82^ on each sequence to generate HMMs on the (http://www.user.gwdg.de/~compbiol/data/hhsuite/databases/hhsuite_dbs/uniprot20_2016_02.tgz) uniprot20_2016_02 database. Second, the HMMER (http://hmmer.org/) program was applied to the set of Uniprot and consensus sequences to search Pfam30 HMM models to generate “HMMER hits”. We searched the Pfam HMMs with the HMMER model of each sequence with the program HHsearch to generate “HH hits”. Last, a greedy algorithm was used to create a unique assignment of a Pfam to each residue in a human sequence, allowing only for short overlaps of < 10 amino acids. We classified Pfam hits into strong or weak hits based on the E-values (i.e. HMMER cutoff = 10^-4^, HH cutoff = 10^-5^) and Pfam HMM coverage (less than 10 residues on N-and C-terminus). We then used greedy algorithm to add hits to the Pfam assignment set in the order of strong HMMER hits, strong HH hits, weak HMMER hits, and weak HH hits with no overlap or less than 10 residues overlap. Finally, we captured additional repeats by adding HMMER weak hits (E-value > 10^-4^) and HH weak hits (E-value > 10^-5^) if there were other assignments within the sequence involving the same Pfam that passed the E-value threshold. The purpose of doing them in this order is to make sure that true hits to the closest Pfam are made first, before hits to other Pfams in the same clan that may score well in the consensus or HHblits hits. If there are two or more assignments of the same Pfam, we checked whether the HMM match states align only once with < 5 residue overlap. If they do, we then combined them into one assignment. We refer to these Pfam assignments as “combined domains”. An assignment was termed a “split domain” if there are > 30 residues between the assigned regions. Those intervening residues are left unassigned or for further assignment. We do not use any Pfam hits with E-value > 10.

### Structure-supported network edge annotation

We identified the structure-supported edges in the interaction network in two stages. First, we utilized the ProtCID database (http://dunbrack2.fccc.edu/ProtCiD/default.aspx) to accumulate all Pfam domain-based interfaces from structures deposited in the Protein Data Bank. Structural domain-based interfaces broadly consist of two categories: domain–domain interfaces and chain–chain interfaces (henceforth referred to simply as Pfam-based and Pfam chain-based interfaces, respectively). In brief, Pfam-based interfaces are defined between Pfam domains, whereas Pfam chain-based interfaces are defined across full-length protein chains (as deposited in the PDB) or “chain architectures”. Pfam chain-based interfaces can be useful in identifying protein interfaces comprising regions of low Pfam coverage that are difficult to capture using domain-based interfaces. Some Pfam domains are shorter than the observed domains in the PDB, and interactions may be recovered by using the full chain sequence. We considered all interfaces larger than 100 Å^2^ found in crystal forms (CFs) of the structures generated from their PDB coordinates.

For Pfam-based interfaces, we extracted 949,296 total interfaces from ProtCID. Some interfaces in the dataset are only found as crystal interfaces but never in the protein biological assemblies and hence could be crystallographic artifacts. We removed them from the dataset and deleted redundant interfaces between the same Pfam domain pairs within a single PDB structure leaving 155,731 interfaces. Approximately 4% of the extracted interfaces contain a “V-set” domain, an Ig-like domain found in diverse protein families including the variable region (F_v_) in antibodies. Although the V-set domains in antibodies directly bind antigens containing diverse domains, the interaction arises from an adaptive immune response and is not likely to be applicable to evolved interactions among proteins in the human proteome. So we omitted the interfaces involving a V-set domain observed in structures containing antibodies (2,538 PDBs)^83^ leading to a final dataset of 12,681 Pfam-based interfaces (2/3 heterodimers, 1/3 homodimers).

Nearly a quarter of these interfaces occur solely within the same chain in PDB structures, we marked such interfaces as “intrachain only” interfaces. Only 54.4% of the Pfam-based interfaces involve a pair of human-specific Pfam domains (domains found at least once in the human proteome) and can be used to identify structure-supported interactions in the human network (Supplementary Fig. S2). Similarly, for Pfam chain-based interfaces, we filtered 423,231 interfaces extracted from PDB structures leading to a final dataset of 4,921 unique Pfam chain-based interfaces. 57.5% of the Pfam chain-based interfaces in the final dataset are between sequences of human-specific Pfam domains and hence are useful to study the human interactome.

Second, we used the human proteome Pfam domain assignments with the curated Pfam-based and Pfam chain-based interface datasets to determine the list of structure-supported edges in the interaction network. After assigning Pfam domains to all the proteins in the interaction network, for each interacting protein pair across the network edge, we evaluated all possible Pfam-based interfaces (*I_edge_*) across the partners as

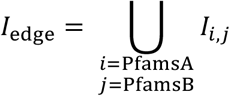

where *I_i,j_* is an interface between any Pfam domains found in interacting proteins A (PfamsA) and B (PfamsB). For example, if protein A contains 3 different Pfams, and protein B contains 4 different Pfams, then *I_edge_* will contain 12 interfaces, not all of which will be represented in the PDB. We then calculated the number of the Pfam-based interfaces (*N_StrI_*) in *I_edge_* that were part of the accumulated dataset of structurally-known Pfam-based interfaces in the PDB (*I*_all_) as

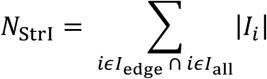

If *N*_StrI_ ≥ 1, we considered the interaction edge to be structure-supported based on an existing Pfam-based interface.

To identify structure-supported interactions in the network that are covered by Pfam chain-based interfaces, we first calculated the pair of chain architectures (*C*_edge_) for the proteins involved in each interaction as

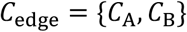

where *C*_A_, *C*_B_ are defined as 〈*i* | *i* ϵ PfamsA〉, 〈*j* | *j* ϵ PfamsB〉 and represent the sequence of Pfam domains in the interacting proteins A, B containing domains PfamsA, PfamsB, respectively. We then calculated the number of interfaces (*N*_StrC_) from the assembled dataset of Pfam chain-based interfaces which support the interaction between the chain architectures *C*_A_, *C*_B_ as

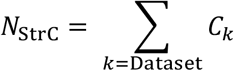

where *C_k_* is defined as

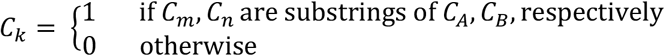

*C_m_, C_n_* are the chain architectures of the interacting protein chains *m, n*, respectively in the domain chain-based interface dataset. *C*_A_ is considered a substring of *C*_A_ if *C*_A_ = *i*_1_ … *i*_PfamsA_ and *C_m_* = *i*_1+*x*_ … *i_m+x_*, where 0 ≤ *x* and *m* + *x* ≤ PfamsA.

If *N*_StrC_ ≥ 1, we considered the interaction edge to be structure-supported based on an existing Pfam chain-based interface.

Finally, we gathered a list of all co-crystallized human protein pairs sharing direct structural interfaces in the PDB-deposited biological assemblies. The list contains a total 3581 PPI from 3108 distinct proteins, including 2086 homodimeric and 1495 heterodimeric interactions. We identified and annotated (“InPDB”) all network edges representing an interaction from this list of interacting proteins.

### Clustering and disease enrichment calculation

We used a three-step process to identify disease-enriched regions of the interaction network.

i. We compiled a list of human disease-associated genes from the MetaCore knowledge base (Thomson Reuters). The list contains 18293 genes associated with 2644 diseases based on evidence of aberrations in DNA, RNA, and/or protein levels.
ii. We divided the interaction network into smaller subgraphs of interacting protein modules using the Markov Cluster Algorithm (MCL).^29^ In MCL, the inflation parameter (*I*) is the primary variable determining the granularity of clustering. To determine the optimal inflation parameter for the study, we iterated over several *I* values ranging from 1.2 to 5 (Supplementary Table S9). For each *I* value, we first clustered the structure-supported network into subgraphs. We next identified large subgraphs with 30 or more proteins and divided them further into smaller subgraphs by clustering using a higher *I* value (*I*=*I*+0.2). We again checked for large clusters and repeated the clustering process on them at increasing *I* values (in steps of 0.2) until clustering did not return any large protein clusters. We used a constant weight (1.0) for all the network edges during clustering.
iii. We finally calculated the percentage of disease-enriched clusters among the MCL-generated clusters generated in the previous step for each *I* value. After annotating individual proteins in the clusters with the associated diseases, we evaluated *P* values for cluster–disease associations using the hypergeometric test (background of 21738 human genes). We adjusted the *P* values for multiple hypothesis testing (Benjamini and Hochberg^84^), and used a 5% FDR cutoff (*P* < 0.05) to identify the disease-enriched protein interaction clusters that were statistically significant. We picked an *I* value of 1.6 for the study as it produced the largest percentage (39.2%) of disease-enriched clusters.

### Disease-causing mutation mapping

We extracted the complete list of literature reported pathological disease-causing mutations from QIAGEN’s Human Gene Mutation Database^47^ (HGMD^®^ 2017.2). We removed mutations 1) that map to intergenic DNA, 2) with multiple observed base pair changes that result in different codons, 3) at transcription initiation sites, 4) in intervening sequences (introns), and 5) before/after initiation/termination codons. This resulted in a final dataset containing 38264 records that captured 5652 unique protein–disease associations between 2445 proteins and 3885 diseases. Nearly 10% of the total records are insertions/deletions and the rest point mutations. We mapped the mutations to regions within proteins and defined all insertions/deletions leading to a shift in the open reading frame and mutations leading to a stop codon as structurally terminal and assumed they would affect all domains C-terminal of the mutation. We considered all other mutations structurally non-terminal affecting only the domain they map to. We then identified all PPI based on domain–domain interfaces involving domains affected by disease-causing mutations, and considered such PPI to be affected by the mutation. We finally analyzed the MCL-generated PPI clusters enriched for disease-associated proteins with known mutations.

### Intrinsically Disordered Region (IDR) prediction

We used disorder predictions in the MobiDB database (http://mobidb.bio.unipd.it/) to estimate the intrinsically disordered regions (IDRs) in the interacting proteins in the network. MobiDB’s intrinsic disorder annotations cover over 80 million UniProt sequences and uses disorder data from diverse sources such as DisProt,^85^ PDB, and ten popular disorder predictors including ESpritz,^80^ IUPred,^87^ DisEMBL,^88^ GlobPlot,^89^ VSL2b,^90^ and JRONN.^91^ We used the MobiDB consensus disorder predictions to calculate the fraction of disordered residues per protein (*fDis*) in 20187 human UniProt sequences.

### Enzyme-substrate interaction extraction

To build a dataset of known enzyme-substrate interactions, we first extracted the list of all modified residues reported in proteins in the UniProt database.^27^ The database contains 4210 protein residue modifications primarily comprising phosphorylation (93.2%), methylation (3.5%), and acetylation (2.3%) events (Supplementary Table S5). We next extracted the annotated enzymes catalyzing the residue modifications found in each substrate protein to identify 2368 unique enzyme–substrate interactions. The modifying enzyme is not known for all modification sites. The interaction dataset is dominated by interactions of kinases (93.5%), as well as containing methyltransferases (3.8%), and acetyltransferases (1.9%) with their substrate proteins.

### Peptide-binding domain identification

We followed a procedure described recently.^24^ Any protein chains with length less than 30 amino acids are defined as peptides. A Pfam–peptide interface is defined with at least 10 C_β_-C_β_ distances < 12 Å, or ≥ 5 atomic contacts with distance < 5 Å. If a peptide is contacting several chains in a biological assembly, the interface with more than 75% atomic contacts is used as the Pfam–peptide interface. For each pair of Pfam–peptide interfaces in a Pfam, the number of Pfam HMM positions shared by the contacts of two interfaces are counted as *N_hmm_*. RMSDs of peptides (*RMSD_pep_*) are calculated after superposing the domain structures. We used the pair_fit routine in PyMOL to align the domain structures via their Pfam HMM positions. We then calculated the minimum RMSD by applying linear least-squares fit on the Pfam interacting regions of two peptides. We clustered Pfam-peptide interfaces using a hierarchical average linkage clustering algorithm by *N*_hmm_ and *RMSD*_pep_. In this method, each interface is initialized to be a cluster. At each step, the two clusters with the highest *N_hmm_* are merged if *N*_hmm_ ≥ 3 and *RMSD*_pep_ ≤ 10 Å.

We next used the assembled dataset of Pfam–peptide interfaces to identify peptide-binding domains in the human interactome. The dataset comprised 516 different Pfam domains found in various domain– peptide interfaces. We calculated the number of interactome PPI involving proteins containing each of the domains and ignored domains found in less than 100 PPI. We also ignored all the enzymatic domains as enzyme–substrate interactions are underrepresented in the human interactome used for this study. We visually inspected the structures of the remaining Pfam–peptide interface clusters to identify the commonly used peptide-binding domains. We finally included a few additional peptide-binding domains found in less than 100 PPI as they were from the same Pfam clans as the commonly used peptide-binding domains.

## Acknowledgments

R.D. acknowledges support from NIH Grant R35 GM122517.

## Contributions

K.P.K., A.L., and R.D. conceived and designed the project. K.P.K., Q.X., and A.L. performed the analyses. K.P.K. and A.L. wrote the paper, and all authors contributed to discussions and editing the paper.

