## Supplementary Data for "Protein domain-based structural interfaces help interpret biologically-relevant interactions in the human interaction network"

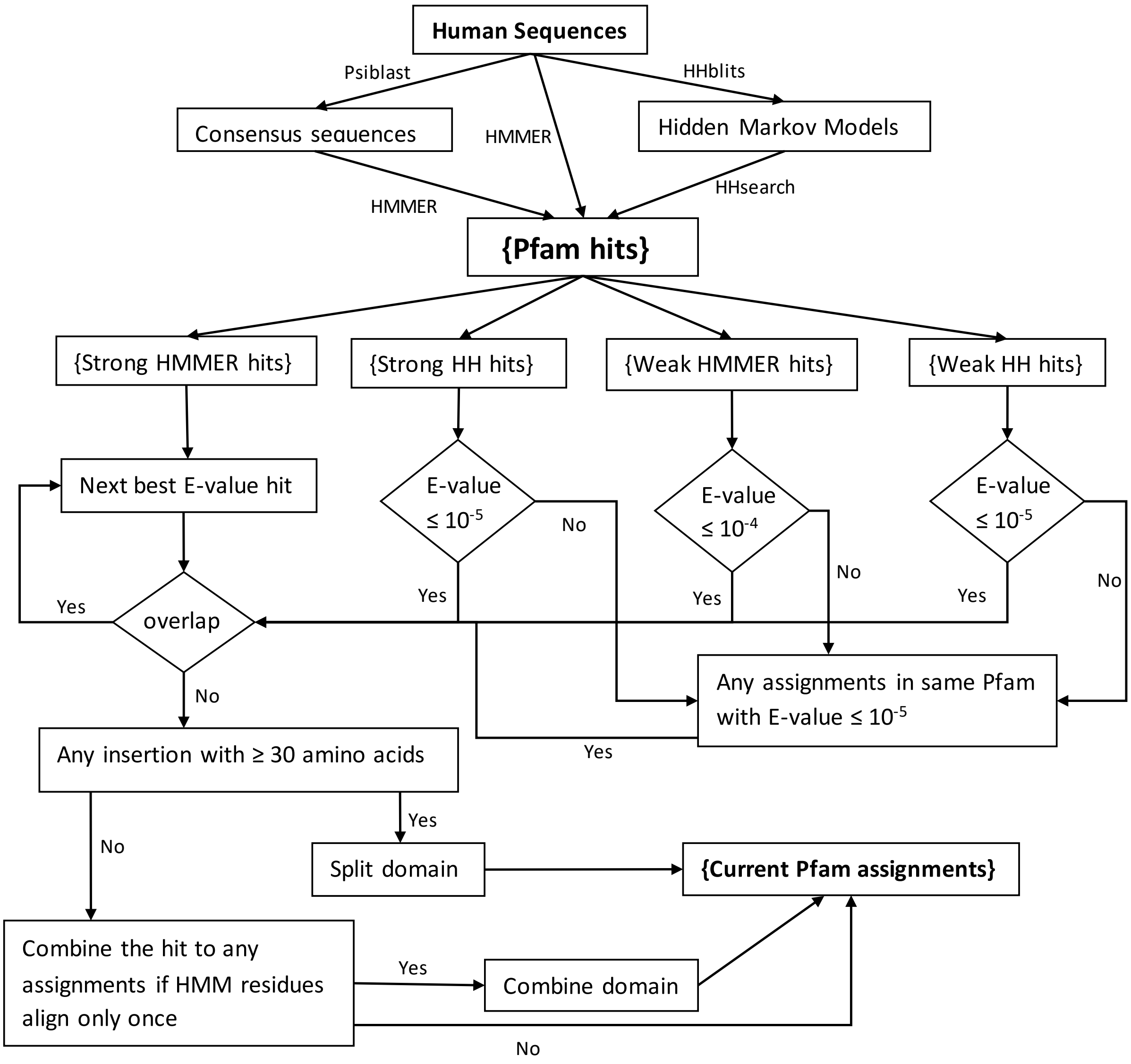


**Supplementary Figure S1:** Flowchart of Pfam assignments to the human proteome using a new greedy algorithm (HuPfam).

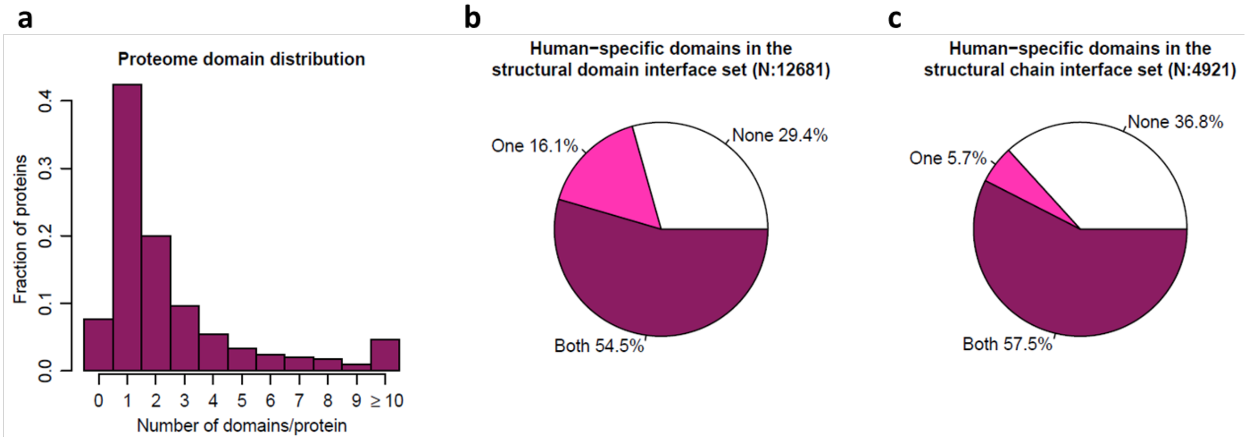


**Supplementary Figure S2:** Human Pfam domains and domain-based interfaces. **a** Distribution of human proteins based on the number of Pfam domains. **b** Human-specific Pfam domain–based interfaces in the dataset accumulated for the study. Interfaces are classified based on whether both, one, or neither of the interacting domains in domain–domain interfaces are found in human proteins. **c** Human-specific Pfam chain-based interfaces in the dataset accumulated for the study. Interfaces are classified based on whether both, one, or neither of the interacting chains contain domains found in human proteins.

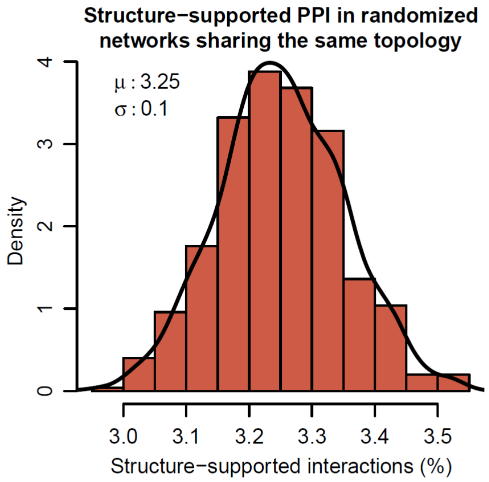


**Supplementary Figure S3:** Structure-supported interactions in randomized PPI networks. Distribution of the percentage of structure-supported interactions across 500 PPI networks generated by randomizing nodes in the human interactome. The node degrees in the networks are preserved during randomization, so all the networks have the same topology as the human interactome. The black curve represents the kernel density estimate of the distribution.

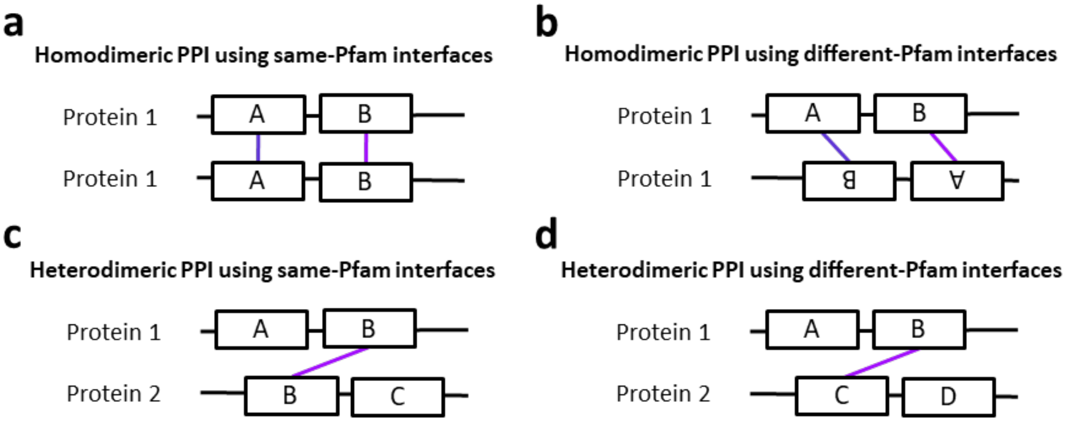


**Supplementary Figure S4:** Domain–domain interfaces used in structure-supported homodimeric and heterodimeric PPI. **a-d** Structure-supported domain–domain interfaces (magenta) between the domains in the interacting proteins (rectangles) are highlighted for different potential PPI modes. Homodimeric PPI are homodimers at the protein-level but could involve interfaces between different Pfams at the domain-level (panel **b**). Similarly, heterodimeric PPI are heterodimers at the protein-level but could be formed by interfaces between same Pfams at the domain-level (panel **c**).

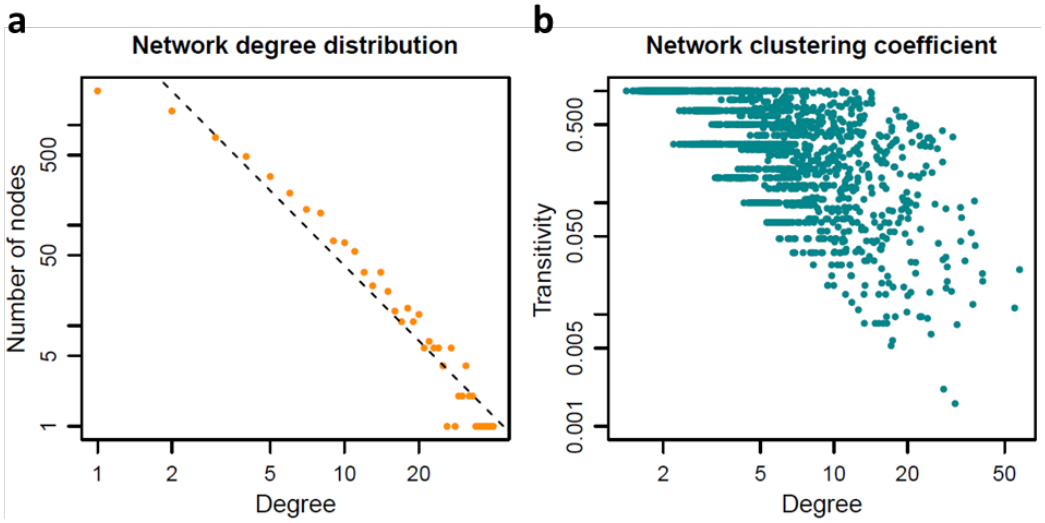


**Supplementary Figure S5:** Topological characterization of the structure-supported network. **a** Degree distribution of the nodes in the structure-supported network obeys the power law illustrating their scale-free topology. **b** Transitivity of the nodes in the structure-supported network gradually drops with an increase in the number of protein interactions indicating a hierarchical network organization.

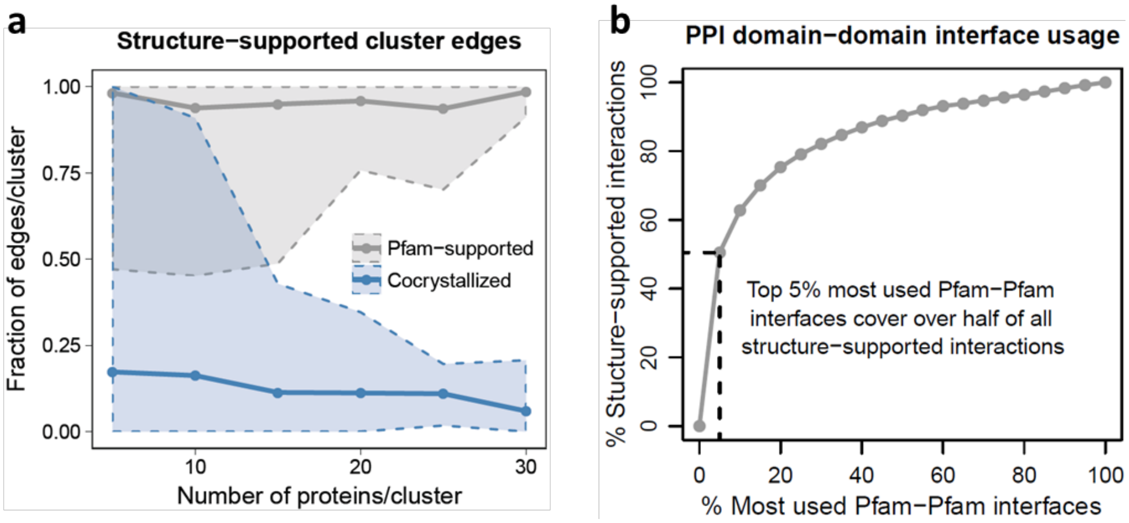


**Supplementary Figure S6:** Structure-supported PPI in interaction clusters and prevalence of domain-based interfaces. **a** Average fractions of Pfam-supported (gray) and co-crystallized (blue) PPI among all the PPI in human interactome clusters of various sizes. The envelopes represent the minimum and maximum edge fractions for each protein cluster size. **b** Cumulative fraction of structure-supported PPI based on the most frequently used domain–domain interfaces.

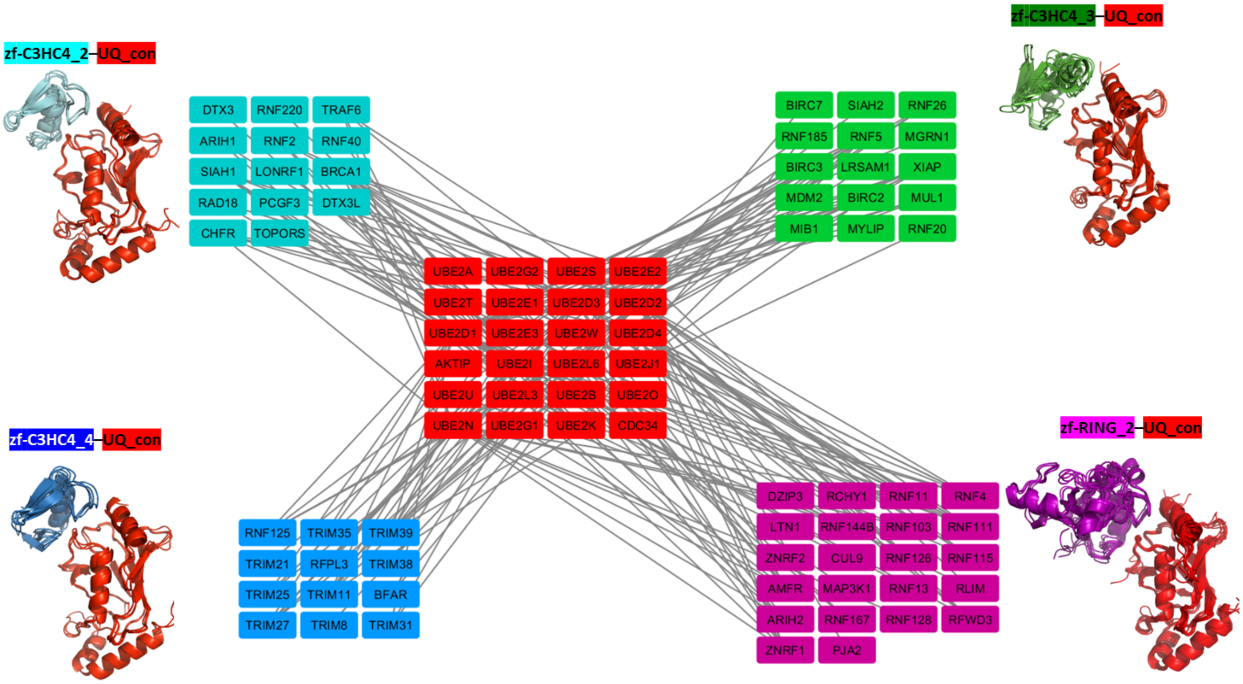


**Supplementary Figure S7:** The protein ubiquitylation PPI network. Structure-supported PPI between E2 ubiquitin-conjugating enzymes (red) and E3 ubiquitin ligases (blue, green, cyan, purple) based on conserved domain–domain interfaces. Proteins are colored based on the domains they contribute to the domain–domain interfaces mediating the PPI.

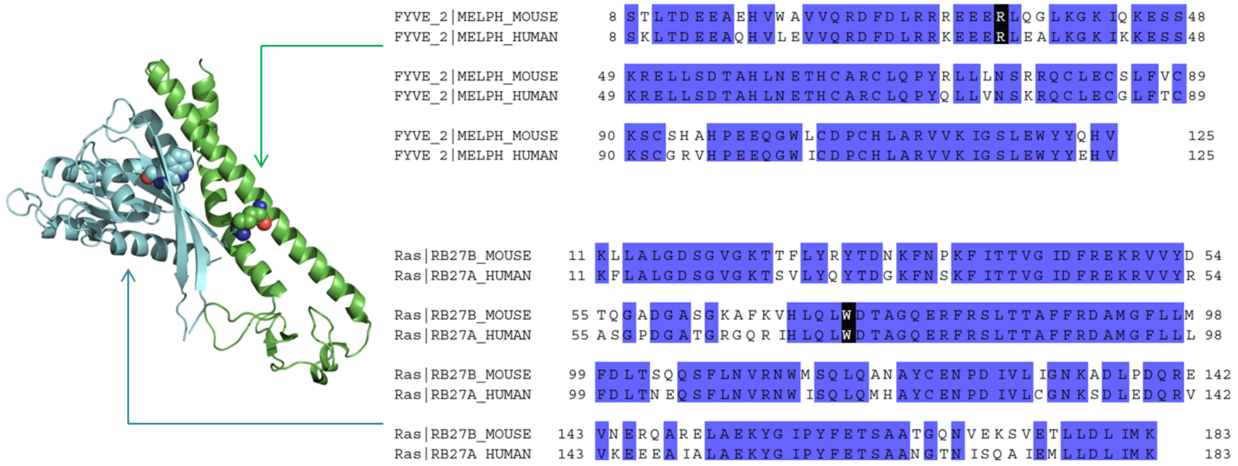


**Supplementary Figure S8:** Griscelli syndrome missense mutations. Ras–FYVE_2 interface in the mouse Rab27B–Slac2-a complex. Sequence alignments demonstrate the high percentage identity between the Ras domains in mouse Rab27B and human Rab27A (~80%), and FYVE_2 domains in mouse Slac2-a and human Slac2-a (85%). Two mutations that abrogate the Rab27A–Slac2-a interaction – Rab27A W73G missense mutation found in Griscelli syndrome type 2 patients, and Slac2-a R35W missense mutation found in Griscelli syndrome type 3 patients are highlighted in black. Residues in the mouse Rab27B–Slac2-a complex homologous to the human Griscelli syndrome-associated mutations are shown in spheres.

**Supplementary Table S1:** Comparison of the Pfam assignments generated by the new greedy algorithm (HuPfam) to existing human proteome Pfam v30 assignments (#Sequences = 18389).

| | #Pfams | #HMM Residues | #Residues | #Splits | #Repeats | #Overlap Residues |
|---|---|---|---|---|---|---|
| HuPfam | 6198 | 5552226 | 5743511 | 988 | 5780 | 25400 |
| Pfam | 6081 | 4976761 | 5246286 | - | 3617 | 33648 |

**Supplementary Table S2:** Pfam domain assignments of UBA1 and CNDP1 human proteins in Pfam and HuPfam. The rows of different split domains are marked in either magenta or orange. In Pfam itself, the E1_FCCH, E1_4HB and UBA_e1_thiolCys domains of protein UBA1 are completely overlapped by two ThiF domains. The M20_dimer domain is also completely covered by Peptidase_M20 domain, which itself also a split domain with 47 residues insertion.

**UBA1 protein**

| | Pfam | SeqBeg | SeqEnd | HmmBeg | HmmEnd | Score | E-value |
|---|---|---|---|---|---|---|---|
| Pfam | ThiF | 55 | 450 | 1 | 240 | 161 | 3.7E-44 |
| | E1_FCCH | 227 | 297 | 1 | 70 | 114 | 3.3E-30 |
| | E1_4HB | 298 | 366 | 1 | 70 | 95 | 2.3E-24 |
| | ThiF | 451 | 952 | 1 | 240 | 248 | 1.2E-70 |
| | UBA_e1_thiolCys | 638 | 884 | 1 | 252 | 326 | 2.3E-94 |
| | E1_UFD | 955 | 1053 | 1 | 93 | 125 | 1.4E-33 |
| FCCC | ThiF | 55 | 225 | 1 | 180 | 161 | 3.5E-47 |
| | E1_FCCH | 227 | 297 | 1 | 70 | 114 | 3.1E-33 |
| | E1_4HB | 298 | 366 | 1 | 70 | 95 | 2.1E-27 |
| | ThiF | 389 | 450 | 181 | 240 | 161 | 3.5E-47 |
| | ThiF | 451 | 633 | 1 | 180 | 248 | 1.1E-73 |
| | UBA_e1_thiolCys | 638 | 884 | 1 | 252 | 326 | 2.0E-97 |
| | ThiF | 885 | 952 | 181 | 240 | 161 | 3.5E-47 |
| | E1_UFD | 955 | 1053 | 1 | 93 | 125 | 1.3E-36 |

**CNDP1 protein**

| | Pfam | SeqBeg | SeqEnd | HmmBeg | HmmEnd | Score | E-value |
|---|---|---|---|---|---|---|---|
| Pfam | Peptidase_M20 | 128 | 500 | 1 | 205 | 115 | 4.6E-30 |
| | M20_dimer | 242 | 400 | 2 | 107 | 46 | 4.4E-09 |
| FCCC | Peptidase_M20 | 128 | 240 | 1 | 115 | 115 | 4.3E-33 |
| | M20_dimer | 241 | 290 | 2 | 48 | 46 | 4.2E-12 |
| | M20_dimer | 337 | 402 | 49 | 107 | 46 | 4.2E-12 |
| | Peptidase_M20 | 406 | 502 | 116 | 205 | 115 | 4.3E-33 |

**Supplementary Table S3:** Dependence of structure-supported interactions on the source used for domain–domain interfaces.

| Pfam–Pfam interface source | BioPlex+ Structure | Y2H+ Structure | Literature+ Structure | Merged+ Structure |
|---|---|---|---|---|
| All protein complexes | 9.8% | 9.3% | 37.9% | 12.6% |
| Exclude co-crystallized human protein complexes captured in interactome PPI | 9.0% | 8.8% | 33.4% | 11.4% |
| Exclude all human protein complexes | 8.5% | 7.9% | 30.1% | 10.5% |

**Supplementary Table S4:** Top connected proteins with more than 30 direct interaction partners in the structure-supported interactome.

| Protein | Degree (Interaction network) | Degree (Structure-supported network) |
|---|---|---|
| GRB2 | 137 | 61 |
| HSP90AB1 | 105 | 52 |
| UBE2D2 | 54 | 42 |
| CUL3 | 67 | 40 |
| ELAVL2 | 98 | 39 |
| NCK2 | 104 | 38 |
| SNRNP70 | 93 | 37 |
| SKP1 | 58 | 36 |
| UBE2D1 | 45 | 36 |
| EGFR | 64 | 32 |
| PIK3R1 | 53 | 32 |
| CDC42 | 37 | 31 |
| CUL5 | 46 | 31 |
| SREK1 | 58 | 31 |
| UBE2D3 | 37 | 31 |
| GDI1 | 33 | 30 |
| UBB | 124 | 30 |

**Supplementary Table S5:** Top enriched GO biological process (BP) terms for the densely-connected proteins (>15 interaction partners) in the structure-supported network.

| GO Biological Process | Sample | Background | Adj. p value |
|---|---|---|---|
| innate immune response | 36/142 | 644/21738 | 1.43E-20 |
| G1/S transition of mitotic cell cycle | 19/142 | 152/21738 | 1.21E-16 |
| Fc-epsilon receptor signaling pathway | 20/142 | 178/21738 | 1.21E-16 |
| epidermal growth factor receptor signaling pathway | 20/142 | 191/21738 | 3.03E-16 |
| regulation of ubiquitin-protein ligase activity involved in mitotic cell cycle | 15/142 | 75/21738 | 3.03E-16 |
| anaphase-promoting complex-dependent proteasomal ubiquitin-dependent protein catabolic process | 15/142 | 80/21738 | 7.14E-16 |
| Fc-gamma receptor signaling pathway involved in phagocytosis | 15/142 | 85/21738 | 1.61E-15 |
| fibroblast growth factor receptor signaling pathway | 18/142 | 160/21738 | 2.84E-15 |
| positive regulation of ubiquitin-protein ligase activity involved in mitotic cell cycle | 14/142 | 71/21738 | 3.06E-15 |
| neurotrophin TRK receptor signaling pathway | 21/142 | 274/21738 | 1.17E-14 |
| axon guidance | 22/142 | 334/21738 | 4.59E-14 |
| mitotic cell cycle | 23/142 | 397/21738 | 1.38E-13 |
| viral process | 26/142 | 544/21738 | 1.51E-13 |
| peptidyl-tyrosine phosphorylation | 15/142 | 134/21738 | 9.36E-13 |
| protein polyubiquitination | 14/142 | 118/21738 | 3.07E-12 |

**Supplementary Table S6:** Top enriched GO molecular function (MF) terms for the densely-connected proteins (>15 interaction partners) in the structure-supported network.

| GO Molecular Function | Sample | Background | Adj. p value |
|---|---|---|---|
| protein binding | 108/142 | 6160/21738 | 1.11E-29 |
| ATP binding | 41/142 | 1444/21738 | 4.95E-14 |
| protein tyrosine kinase activity | 13/142 | 79/21738 | 3.79E-13 |
| cyclin-dependent protein serine/threonine kinase activity | 9/142 | 32/21738 | 2.85E-11 |
| threonine-type endopeptidase activity | 7/142 | 21/21738 | 2.60E-09 |
| ubiquitin-protein transferase activity | 15/142 | 266/21738 | 1.10E-08 |
| cyclin binding | 6/142 | 19/21738 | 6.97E-08 |
| SNAP receptor activity | 6/142 | 23/21738 | 2.22E-07 |
| neurotrophin TRKA receptor binding | 4/142 | 5/21738 | 2.66E-07 |
| phosphatidylinositol 3-kinase binding | 5/142 | 19/21738 | 3.30E-06 |
| acid-amino acid ligase activity | 6/142 | 38/21738 | 4.07E-06 |
| transcription coactivator activity | 11/142 | 236/21738 | 1.03E-05 |
| insulin receptor substrate binding | 4/142 | 11/21738 | 1.18E-05 |
| ErbB-3 class receptor binding | 3/142 | 4/21738 | 2.00E-05 |
| insulin binding | 3/142 | 4/21738 | 2.00E-05 |

**Supplementary Table S7:** Enzyme-substrate interactions extracted from the UniProt database.

| Modification type | Number of enzymatic residue modifications | Number of unique enzyme-substrate interactions |
|---|---|---|
| Phosphorylation | 3838 | 2215 |
| Methylation | 144 | 89 |
| Acetylation | 95 | 46 |
| Hydroxylation | 35 | 11 |
| Deimination | 4 | 4 |
| Decarboxylation | 2 | 2 |
| ADP-ribosylation | 2 | 1 |
| **All** | **4120** | **2368** |

**Supplementary Table S8:** PPI mediated by peptide-binding domains. The number of peptide-binding domains for each peptide recognition motif are shown. The number of estimated PPI involving peptide-binding domains and the percentage of such PPI that are also supported by known domain–domain interfaces in the human interactome are also listed. Majority of the PPI involving hub proteins also use peptide-binding domains.

| Peptide recognition motif | Number of peptide-binding domains | Number of PPI | Domain–domain supported PPI | PPI involving hub proteins |
|---|---|---|---|---|
| Post-translationally modified (Methylation, Acetylation, Phosphorylation) | 23 | 4559 | 19.1% | 65.7% |
| Proline-dependent | 9 | 3926 | 20.7% | 64.5% |
| Other | 20 | 5409 | 14.4% | 56.8% |
| All | 52 | 12231 | 15.8% | 61.4% |

**Supplementary Table S9:** Iterations to identify the optimal inflation parameter for MCL clustering.

| I | NCLUS | NCLUS1 | NCLUS2 | NCLUS.DIS | NCLUS.DIS.PER | MODULARITY |
|---|---|---|---|---|---|---|
| 1.2 | 1397 | 341 | 425 | 536 | 38.40% | 0.76 |
| 1.4 | 1424 | 341 | 430 | 552 | 38.80% | 0.76 |
| 1.6 | 1563 | 343 | 472 | 612 | 39.20% | 0.74 |
| 1.8 | 1732 | 363 | 568 | 655 | 37.80% | 0.72 |
| 2 | 1898 | 392 | 691 | 669 | 35.20% | 0.68 |
| 2.2 | 1987 | 418 | 735 | 674 | 33.90% | 0.66 |
| 2.4 | 2074 | 454 | 802 | 664 | 32% | 0.64 |
| 2.6 | 2166 | 501 | 858 | 659 | 30.40% | 0.61 |
| 2.8 | 2254 | 559 | 898 | 646 | 28.70% | 0.59 |
| 3 | 2309 | 604 | 921 | 633 | 27.40% | 0.58 |
| 4 | 2610 | 912 | 964 | 589 | 22.60% | 0.52 |
| 5 | 2797 | 1126 | 969 | 558 | 19.90% | 0.49 |
